# Calcium Signaling and Glycerol-3-Phosphate Metabolism Coordinate Extracellular Vesicle Biogenesis in *Cryptococcus neoformans*

**DOI:** 10.64898/2026.11.25.769125

**Authors:** Alicia C. Piffer, Jean-Yves Coppée, Pierre-Henri Commere, Corinne Maufrais, Frédérique Moyrand, Juliana Rizzo, Cosmin Saveanu, Guilhem Janbon

## Abstract

Extracellular vesicles (EVs) are increasingly recognized as important mediators of fungal physiology and host–pathogen interactions, but the mechanisms governing their biogenesis remain poorly understood. Here, we combined time-course RNA sequencing with genome-wide genetic screening to define the regulatory network controlling EV production in the human fungal pathogen *Cryptococcus neoformans*. Transcriptomic analysis identified an EV-associated regulon enriched in genes involved in vesicular trafficking, cell wall remodeling, lipid metabolism, mitochondrial function and ion homeostasis. Our screen further identified diverse regulators of EV production, highlighting endosomal trafficking, mitochondrial metabolism and calcium homeostasis. Cell viability was only weakly associated with EV production, arguing against cell lysis as the primary mechanism of EV release. Genetic and pharmacological analyses established calcium-dependent calcineurin signalling as a major regulator of EV production. We identified the calcineurin target Had1 as a positive regulator of EV production and demonstrate that Had1 functions as a glycerol-3-phosphate phosphatase. Loss of *HAD1* caused glycerol-3-phosphate accumulation and extensive remodeling of cellular phospholipids, accompanied by a marked reduction in EV production. Together, these findings identify a calcium–calcineurin–Had1 pathway linking intracellular signalling and lipid homeostasis to fungal EV biogenesis.

## Introduction

It is estimated that there are between 2.2 and 3.8 million fungal species, yet only around 200 of these are known to cause human disease^1, 2^. Recent estimates suggest that nearly one billion superficial or invasive fungal infections occur each year, resulting in around 1.6 million deaths worldwide^3^. Beyond their pathogenic potential, fungi also play essential, yet often overlooked, roles within the human microbiota. These organisms, collectively referred to as the mycobiome, interact with bacteria and other microorganisms to maintain microbial homeostasis and modulate host immune responses^4^. Growing evidence indicates that fungi actively shape immune regulation and influence how the immune system discriminates between commensal and pathogenic microbes^5^. Accordingly, disturbances of the fungal community, or dysbiosis, have been associated with various conditions, including inflammatory bowel disease, allergies, cancer, neurodegenerative disorders, and autoimmune diseases^4, 6, 7^. In recognition of their growing impact on public health, the World Health Organization published a list of priority fungal pathogens in 2022, emphasizing the urgent need for research and development in this area^8^.

Extracellular vesicles (EVs) are lipid bilayer nanoparticles produced by organisms across all kingdoms of life ^9^. They have been shown to mediate intra- and inter-kingdom cell-to-cell communication^10^. They have been also proposed as means of promoting cell homeostasis^11^. EVs have been implicated in a wide range of physiological and pathological processes in mammals, as well as in numerous host–pathogen interactions^10^. For instance, EVs from human fungal pathogens can elicit pro- or anti-inflammatory responses in macrophages and monocytes^12^. EVs have also been shown to promote intracellular replication of *Cryptococcus neoformans* ^13^ and to regulate filamentation in *Candida albicans*^14^. They also modulate antifungal resistance in these two major pathogenic yeasts^15,16^. EVs from *Sporothrix* species enhance fungal adhesion to endothelial cells^17^. Proteomic analyses have revealed that human fungal pathogens EVs are enriched in major antigenic components, which is consistent with the observation that immunization with EVs confers protection in models of *C. albicans* and *C. neoformans* infections^18, 19^. In addition to proteins, DNA associated with fungal EVs can regulate inflammatory responses *in vivo* through activation of the cGAS–STING pathway^20, 21^. In plant-pathogenic fungi, EVs also play a key role in plant–pathogen interactions^22^. Overall, while *in vivo* mechanistic data remains limited, these findings support a central role for EVs in host–pathogen interactions in fungal pathogens, analogous to the well-established roles of EVs in bacterial and parasitic infections^23, 24^.

Whereas extensive literature describes the pathways and mechanisms involved in EV biogenesis in mammals^25^, these processes remain comparatively understudied in fungi^26^. In mammalian systems, several genome-wide screens have identified numerous genes that regulate EV production and uptake ^4–17^. In contrast, only a limited number of genes implicated in fungal EV biogenesis have been identified, and the mechanisms by which EVs are released into the extracellular space remain largely hypothetical^26^. In *C. albicans* and *Saccharomyces cerevisiae*, deletion of selected Endosomal Sorting Complex Required for Transport (ESCRT) machinery components alters EV production, supporting a role for the endosomal pathway and the existence of an exosome-like biosynthetic route^15, 27^. Conversely, in *Aspergillus fumigatus*, microscopic observations of EV-like particles budding from the plasma membrane of regenerating protoplasts following cell wall digestion suggest the presence of a microvesicle-like pathway^28^. We recently conducted two genetic screens in *C. neoformans*, targeting transcription factor and kinase mutant collections, respectively, to identify regulators of EV production^16, 29^. The kinase mutant screen identified the cell wall integrity (CWI) pathway as a negative regulator of EV production, highlighting the role of the cell wall in this process and leading to the characterization of a previously unrecognized class of fungal vesicles resembling mammalian apoptotic EVs ^29^. Screening of the transcription factor mutant library identified the HAP complex, a key regulator of mitochondrial function, as a major positive regulator of EV production in *C. neoformans* in agreement with the observed timing of EV production being restricted to the transition between exponential to stationary growth phase^16^.

Time-course RNA-seq has been widely applied across diverse organisms to investigate dynamic changes in gene expression in response to stress, during development, and throughout circadian cycles^30–33^. This powerful approach provides a global view of transcriptomic dynamics over time and enables the identification of novel genes involved in specific biological pathways. Such analyses typically rely on the assumption that genes displaying coordinated temporal expression patterns are likely to participate in shared regulatory networks or functional pathways. This strategy has been also used to study human fungal pathogens. For example, time-resolved gene expression profiling of *C. albicans* during macrophage infection revealed coordinated host–pathogen transcriptional dynamics^34^, while temporal transcriptomic analyses of *A. fumigatus* biofilm development identified stage-specific metabolic adaptations required to maintain biofilm maturation^35^. In *C. neoformans*, time course RNA-seq has been used to study the cell-cycle regulated transcriptional program^36^ and more recently to study cell wall modification upon filamentation induced by glucosamine^37^.

In this study, we employed two complementary approaches to elucidate the regulatory pathways governing EV biogenesis in *C. neoformans*. Time-course RNA sequencing enabled the identification of genes whose expression is dynamically modulated during EV production, thereby defining an EV-associated regulon and the pathways involved. In parallel, a genome-wide screen uncovered key regulators of the EV biosynthetic machinery, which we further characterized using complementary biochemical analyses to elucidate the underlying mechanisms. Our findings identified calcium homeostasis as a central determinant of EV production and revealed a specific role for mitochondrial metabolism and calcineurin signaling in this process. We further demonstrated that Had1, a previously identified calcineurin target, functions as a glycerol-3-phosphate phosphatase. Lipidomic profiling revealed an Had1-dependent modulation of phospholipid metabolism, suggesting a central role for these phospholipids in EV formation. Overall, our study provides the first global view of genes implicated in EV production in *C. neoformans* and offers a comprehensive description of a regulatory pathway controlling EV biogenesis in fungi.

## Results

### Time course RNA-seq analysis of EV production

We previously demonstrated that EV production in *C. neoformans* is regulated by both growth conditions and growth stage^16^. Under our experimental conditions, (i.e. synthetic dextrose, SD at 30 °C), EV release peaked during the transition from exponential to stationary phase, corresponding to 15–20 h post-plating^16^. To investigate the mechanisms underlying EV production, we monitored gene expression dynamics across five time points (14, 16, 18, 20, and 24 h post-plating), using cells from early stationary phase in YPD liquid medium as the reference (time 0)^38^. All experiments were performed in biological triplicate. Time-series RNA-seq data were analyzed using a Dirichlet process Gaussian process mixture model to identify genes with similar temporal expression profiles^39^. This analysis produced 28 co-expression clusters, each containing between 37 and 153 genes (mean = 99 genes), encompassing a total of 2,798 genes (Fig. 1A; Table S1A).

**Figure 1.**
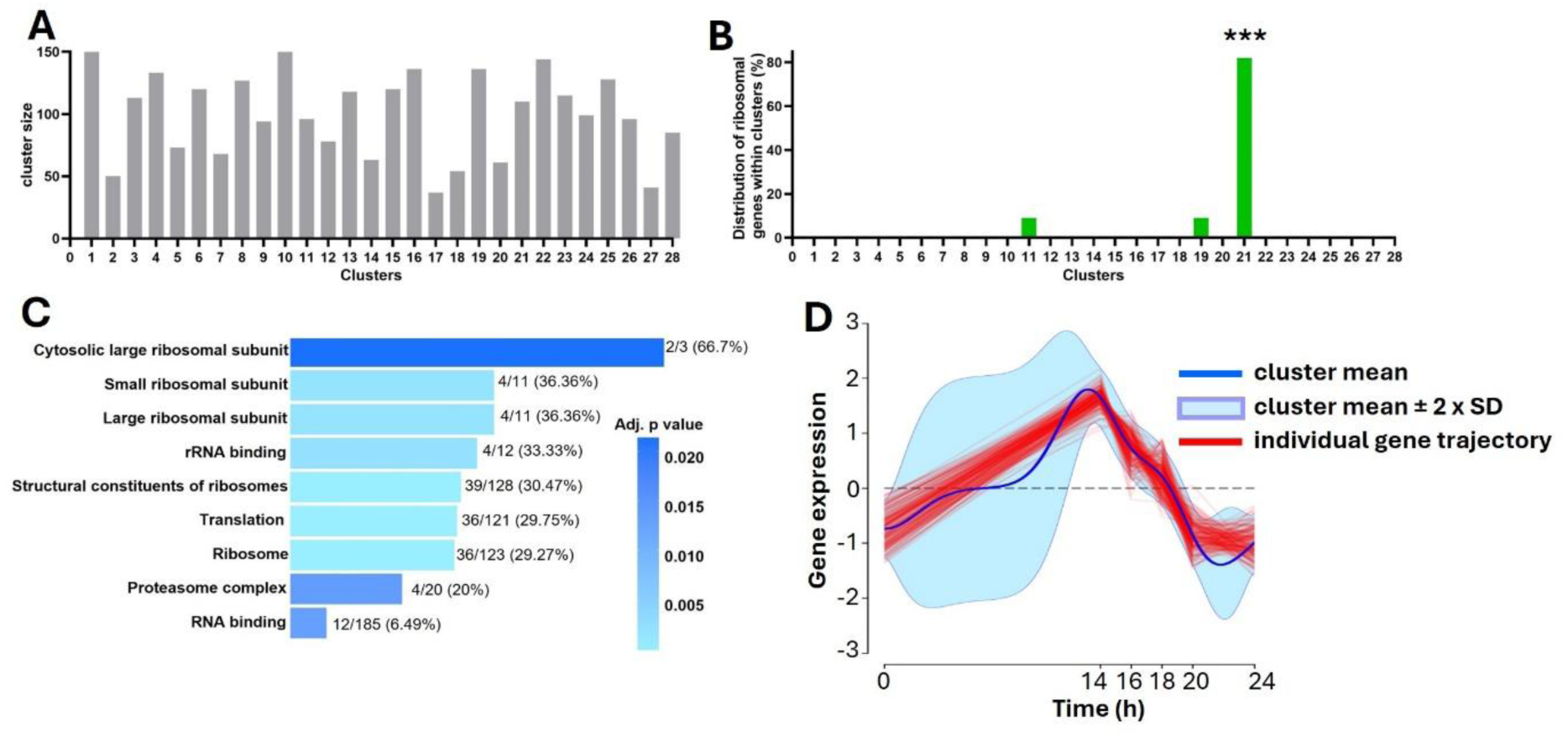
| Clustering of temporal gene expression profiles during the exponential-to-stationary phase transition in *C. neoformans*. A. Sizes of the 28 identified clusters. B. Cluster distribution of ribosomal protein–encoding genes. Statistical significance was assessed using a one-tailed hypergeometric test to evaluate sample enrichment across the 28 clusters, correcting for differences in population sizes. *p*-values were adjusted for multiple testing using the Benjamini-Hochberg FDR procedure. *** indicates adjusted q value < 0.001. C. Significantly enriched Gene Ontology (GO) terms among C21 gene set, as identified using FungiFun3 with default parameters^42^. The percentage indicates the ratio of genes from each cluster associated with a given GO term relative to the total number of background genes in that category. Cluster C21: GO:0022625 (cytosolic large ribosomal subunit); GO:0015935 (small ribosomal subunit); GO:0015934 (large ribosomal subunit); GO:0019843 (rRNA binding); GO: 0003735 (Structural components of ribosomes); GO:0006412 (translation); GO:0005840 (ribosome); GO:0000502 (proteasome complex); GO:0003723 (RNA binding). D. Time-course expression profiles of cluster C21 genes.

To assess the quality of this dataset, we first examined genes encoding ribosomal proteins, reasoning that they should be co-regulated^40^ and therefore grouped into a limited number of clusters. As expected, 82% (n = 41) of all clustered ribosomal protein-encoding genes were assigned to cluster C21, which contained 110 genes (Fig. 1A) (fold enrichment = 36.0, *q-value* < 0.001) (Fig.1B). Consistent with this finding, Gene Ontology terms related to ribosome biogenesis, translation, and RNA binding were significantly enriched in this cluster (Fig. 1C). Finaly, genes within this cluster were strongly downregulated between 16 h and 24 h, reflecting the expected global reduction in protein synthesis machinery as cells enter stationary phase^41^ (Fig. 1D).

To explore transcriptional changes associated with EV production, we looked at the 38 genes encoding EV-associated proteins previously identified in *C. neoformans*^18^. Whereas only 12 of these genes were assigned to co-expression clusters, half of them were grouped in cluster C14 (fold enrichment = 9.2, *q-*value < 0.001), including *MP88* and *VEP1*, which encode the two most abundant EV proteins^18^ (Fig. 2A). Genes within this cluster showed maximal expression during the exponential-to-stationary transition, consistent with a coordinated regulation of EV biogenesis during the transition to stationary phase (Fig. 2B).

**Figure 2.**
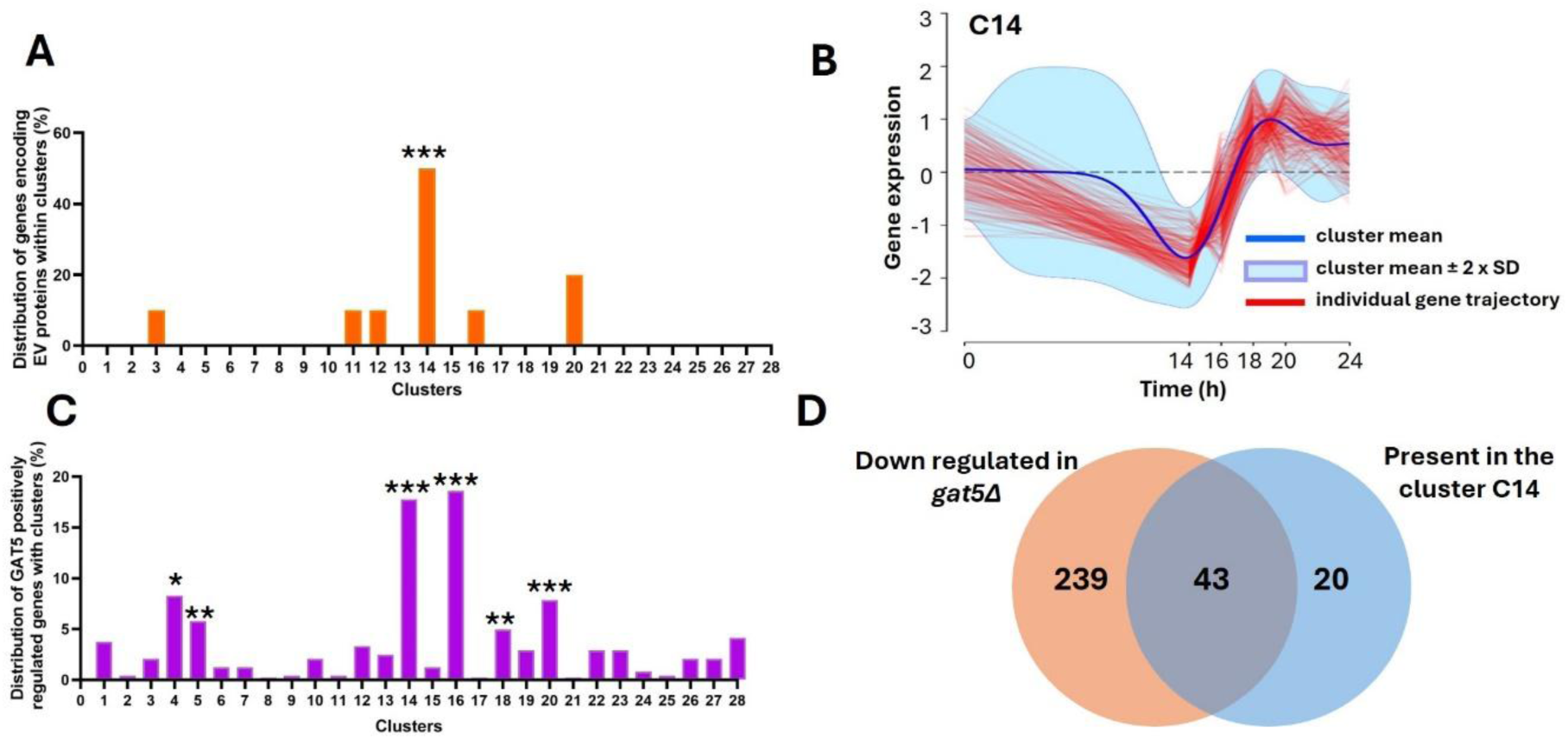
| Genes belonging to cluster 14 are co-regulated with EV production. A. Cluster distribution of EV protein–encoding genes. B. Time-course expression profiles of cluster C14 genes. C. Cluster distribution of *GAT5* positively regulated genes D. Venn diagram showing the enrichment of *GAT5* positively regulated genes within cluster C14. Statistical significance was assessed using a one-tailed hypergeometric test to evaluate sample enrichment across the 28 clusters, correcting for differences in population sizes. *p*-values were adjusted for multiple testing using the Benjamini-Hochberg FDR procedure. *** indicates adjusted *q*-value < 0.001.

We previously identified two transcription factors, Hap2 and Gat5, as positive regulators of EV production in *C. neoformans*^16^. RNA-seq analysis showed that they act within the same regulatory pathway, with Hap2 controlling *GAT5* expression and suggested a direct transcriptional regulation of the genes encoding EV associated proteins^16^. Accordingly, cluster 14 is the most enriched cluster in *GAT5* positively regulated genes (fold enrichment = 65.9, *q*-value < 0.001) (Fig. 2C), and 68% of the cluster C14 genes are downregulated upon *GAT5* deletion (Fig. 2D). Overall, these results suggest that the 63 genes comprising cluster 14 are likely to constitute a regulon associated with EV production in *C. neoformans*.

We used manual gene annotation and Gene Ontology enrichment analysis to better describe this EV regulon (Table S1B). We observed a strong representation of polysaccharide-modifying enzymes (*KRE6, KRE61, CHS8, GAS1, SPR1, CPS1, ROT2, CDA3, AMS1,* and *CWH43*), many of which are known EV cargo components (Fig.3). This finding supports the idea that cell wall remodeling enzymes may participate in EV biosynthesis and/or export pathways as previously suggested^26^. The presence of the Cell Integrity Pathway kinase Bck1 encoding gene in this cluster further supports this hypothesis and is in good agreement with our recent data^29, 43^. Cluster C14 also included genes involved in ergosterol metabolism and its regulation (*ERG11, ERG3,* and *SRE1*), consistent with the reported co-regulation of EV production and fluconazole (FLC) resistance^16, 44, 45^. Notably, numerous genes in this cluster are implicated in vesicle composition^18^ and trafficking pathways (Table S1B). For example, *MIL1* in *S. cerevisiae* encodes a predicted lipase that binds the AP-1 protein Amp2, a key mediator of cargo sorting in clathrin-coated vesicles^46^, whereas *APT2* encodes an homolog of the *S. cerevisiae* lipid flippase Dnf1, which mediates phospholipid transport across membranes, an activity essential for vesicle budding and membrane trafficking^47^. Flippases and their regulatory element *Cdc50* have also previously been shown to regulation membrane asymmetry, EV production and protein cargo composition in *C. neoformans*^48, 49^, consistent with the inclusion of *APT2* in this EV regulon.

**Figure 3.**
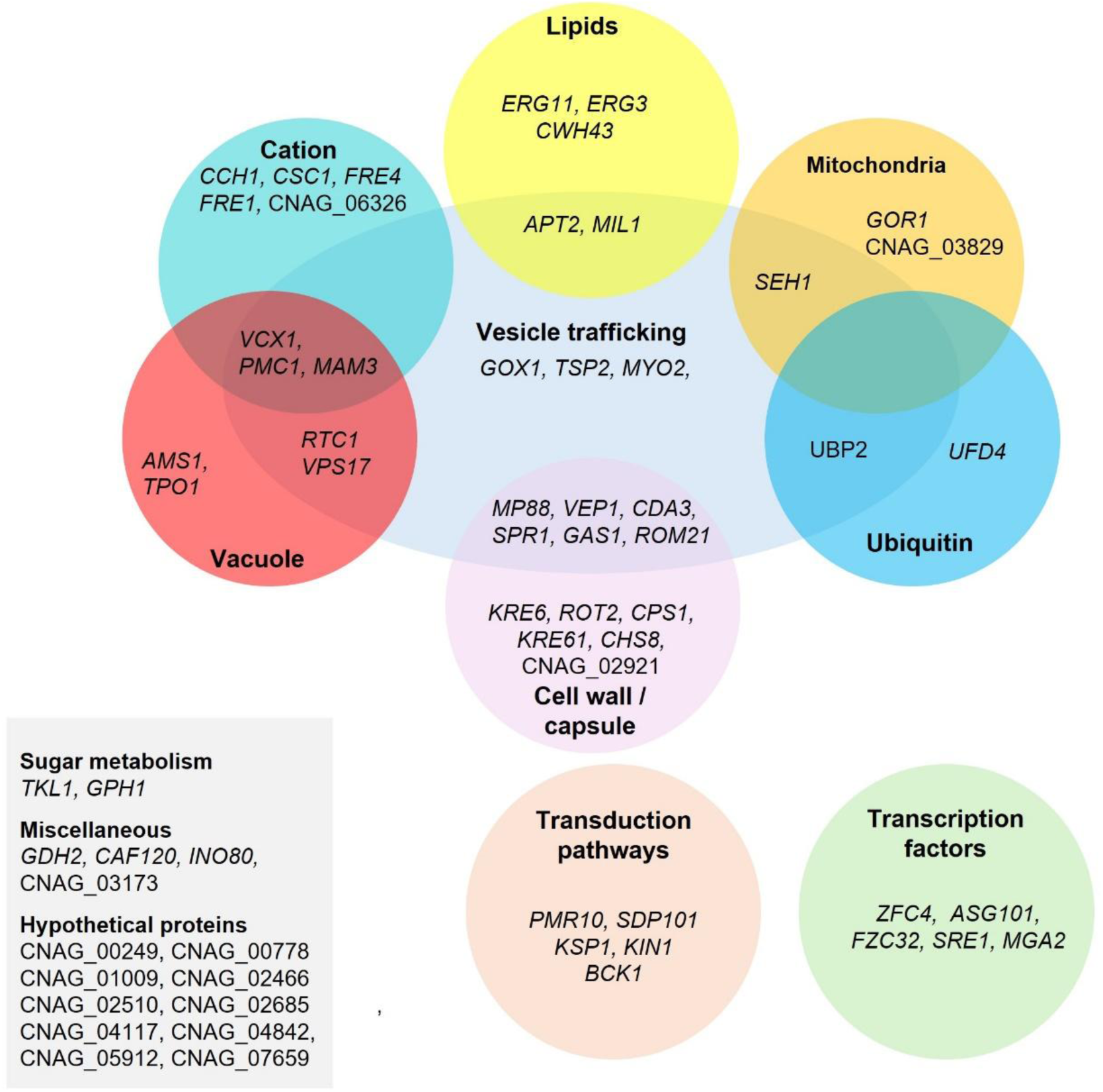
| The EV regulon. Functional annotation of the 63 genes comprising cluster C14 revealed an enrichment for genes coding for vesicle trafficking and related pathways. Gene names correspond to the most significant homologs in *S. cerevisiae*, unless a *C. neoformans* gene designation was already established. Genes were grouped into categories based on their predicted or experimentally validated functions.

Finaly, we noted the presence of several genes implicated in iron, magnesium and calcium homeostasis (*VCX1, PMC1, CCH1, CSC1, MAM3, FRE1,* and *FRE4*) associated with the EV regulon^50, 51^. Consistent with this observation, several GO-terms linked to calcium and monatomic ions homeostasis were enriched in cluster C14 indicating that cations may play a role in regulating EV production (Fig. 4A). Also consistent with this observation, we noted the presence of several proteins implicated in vacuole and mitochondria metabolism, with both organelles known to participate in intracellular calcium concentration regulation^52, 53^. Calcium ion homeostasis and the calcineurin pathway have been already reported to play major roles in the regulations of vesicle trafficking in different organisms including in fungi^54^. For instance, most calcineurin pathway mutant strains were reported to be also deficient in different aspects of vesicular trafficking in *Candida glabrata* and *Schizosaccharomyces pombe*^55, 56^. Consistently, analysis of the cluster distribution of genes regulated by the calcineurin pathway in *C. neoformans*^57^ revealed that cluster C14 was the most enriched (fold enrichment = 15.3; q value < 0.001) (Fig. 4B).

**Figure 4.**
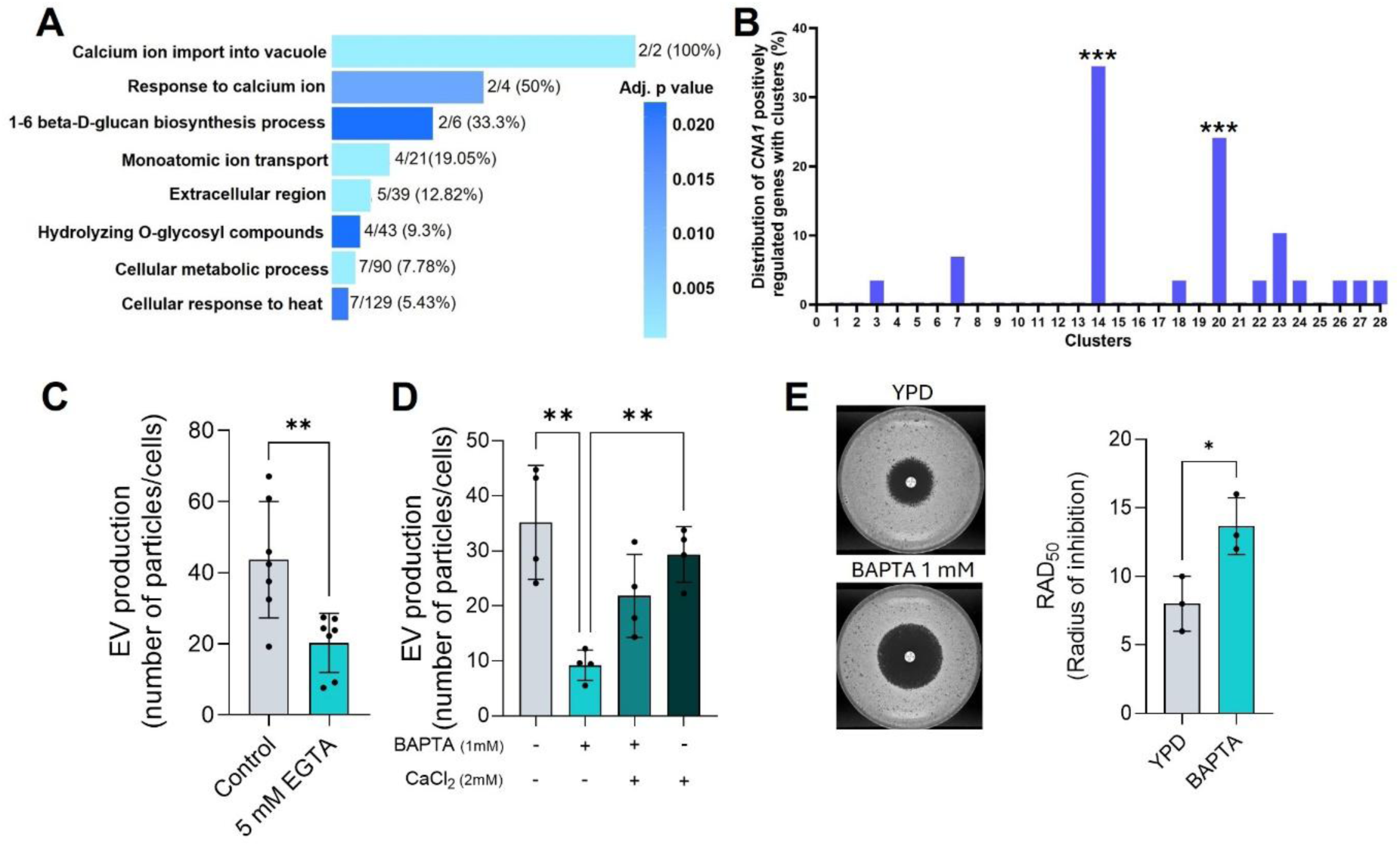
| Calcium homeostasis regulates EV production and FLC resistance. A. Significantly enriched Gene Ontology (GO) terms among the C14 gene set, as identified using FungiFun3 with default parameters^42^, The percentage indicates the ratio of genes associated with a given GO term relative to the total number of background genes in that category. GO 0140146: calcium ion import into vacuole; GO 0051592: response to calcium ion; 0006078; 1-6 beta-D-glucan biosynthesis process; 0006811: monoatomic ion transport; 0005576 : Extracellular region; 0004553: hydrolase activity, hydrolyzing O-glycosyl compounds; GO 0005975: Cellular metabolic process GO 0034605: cellular response to heat. B. Cluster distribution of *CNA1* positively regulated genes^57^. Statistical significance was assessed using a one-tailed hypergeometric test to evaluate sample enrichment across the 28 clusters, correcting for differences in population sizes. p-values were adjusted for multiple testing using the Benjamini-Hochberg FDR procedure. *** indicates adjusted *q*-value < 0.001. C. EV production in the presence of the calcium chelator EGTA. ** means *p* < 0.01 (unpaired Student’s *t*-test). EVs were isolated using the quick protocol and their concentration evaluated by NanoFCM. D. EV quantification in the presence of BAPTA (calcium chelator), CaCl₂, or both. ** means *p* < 0.01 (ANOVA followed by Tukey’s multiple comparison test). EVs were isolated using the quick protocol and their concentration evaluated by NanoFCM. E. Representative FLC susceptibility assay by disk diffusion (right) and RAD50 values (cm) determined using diskImageR (left). Error bars represent mean ± SD. * means *p* < 0.05 (unpaired Student’s *t*-test).

To directly assess the effect of calcium on EV release, we examined the impact of adding broad-spectrum (EGTA) and calcium-specific (BAPTA) chelators to the culture medium. As shown in Fig. 4C,D, cation depletion markedly reduced EV production, an effect that was reversed by cation supplementation. Interestingly, as previously observed in *Aspergillus fumigatus* BAPTA treatment also altered FLC susceptibility (Fig. 4E), further linking EV biogenesis, antifungal response, and calcium homeostasis^58^. Together, these findings confirm the phase-dependent regulation of EV production in *C. neoformans* and highlight calcium signaling as a central regulatory mechanism, paralleling reports from mammalian EV systems^59–61^.

### Genome-wide screening for EV production regulators in *C. neoformans*

In parallel with the time-course RNA-seq analysis, we performed a genome-wide screen to identify key regulators of EV production. We previously developed and applied a rapid EV isolation protocol to screen two mutant libraries (one of 150 transcription factor mutant strains and another of 130 kinase mutant strains of *C. neoformans*) to identify regulators of EV production ^16, 29, 62^. Although this protocol is faster than the classical ultracentrifugation-based approach, it remains laborious and unsuitable for large-scale analyses such as the genome-wide mutant collection recently generated by the Madhani laboratory at UCSF which comprises 4,401 mutants ^63^. To overcome this limitation, we leveraged the previously observed relationship between EV production and FLC susceptibility^16, 29^. We reasoned that a pre-screening step based on FLC susceptibility could serve as an enrichment strategy, thereby reducing the number of mutants to be tested for EV production. We therefore first assessed the FLC susceptibility of all 4,401 strains by spotting them onto YPD medium containing three concentrations of FLC (8, 16, and 32 µg mL⁻¹). Visual inspection identified 310 mutant strains with altered growth compared to control plates lacking FLC. Twenty-three strains unable to grow or severely impaired on SD medium were excluded from further analyses. We next quantified EV production in each of the 287 remaining strains using a NanoFCM (NanoFCM, China) instrument (Table S2). Based on natural variability among wild-type (WT) samples, thresholds corresponding to 70% and 130% of WT EV concentrations were defined to identify mutants with reduced or elevated EV production, respectively. Using these criteria, we identified 158 genes encoding for regulators of EV production (Fig.5A, B; Table S2).

**Figure 5.**
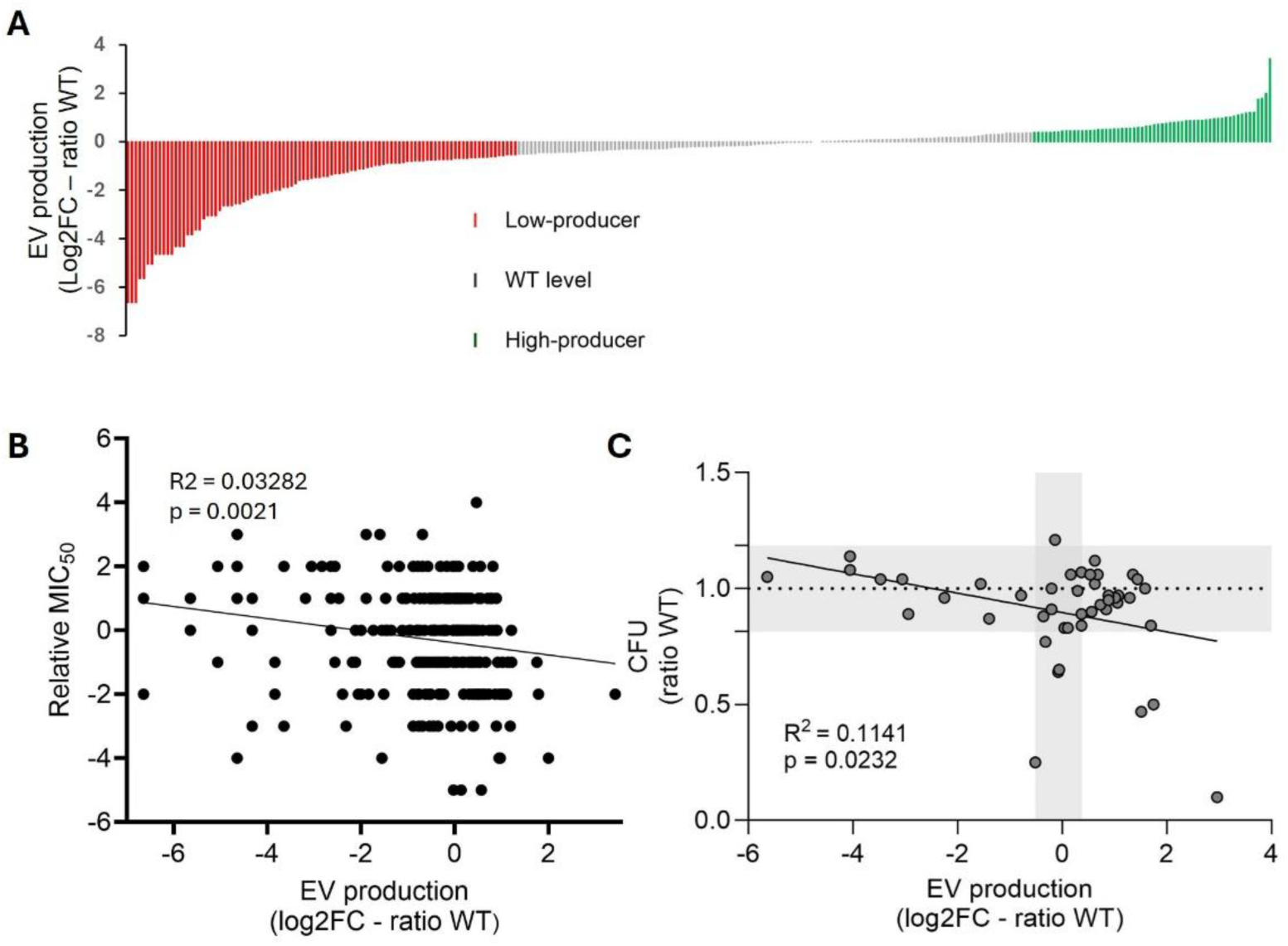
| Genome-wide screening for EV production mutants. **A.** Quantification of EV production by NanoFCM in 287 FLC-altered susceptibility mutant strains from the Madhani collection. Data are presented as log₂ fold change relative to the WT. Mutants producing less than 0.7-fold the EV level of the WT are shown in red; those producing at least 1.3-fold more EVs than the WT are shown in green. **B.** Correlation between FLC susceptibility (measured by MIC₅₀) and EV production (log₂ fold change relative to the WT) as evaluated using a NanoFCM instrument. **C.** Correlation between mutant viability (measured by CFU counts) and EV production (log₂ fold change relative to the WT) as evaluated using a NanoFCM in triplicate. Grey areas indicate the range of variability observed in WT samples.

Consistent with our initial hypothesis, 60% of FLC-sensitive mutants and 51% of FLC-resistant mutants exhibited altered EV production. To definitively prove the validity of our strategy, we randomly selected 41 additional mutants from the Madhani collection and tested them for EV production. Only three of them displayed altered EV production (Table S3; Fig. S1). Follow-up spot assays revealed that one of them was FLC-sensitive and had been missed in the initial screen (Fig. S1). Finaly, to examine whether EV production levels correlated with antifungal susceptibility, we measured the MIC₅₀ values of each mutant in liquid YPD medium and plotted them against EV production values obtained by NanoFCM (Fig. 5B). Although statistically significant anticorrelation was observed between FLC resistance and EV production, this was modest and the MIC_50_ value of given strain is not predictive of its EV production phenotype. Collectively, these analyses validate our screening strategy and reinforce the link between EV production and FLC susceptibility in *C. neoformans*.

### Cell viability is not a reliable predictor of EV release

Several mechanisms have been proposed to account for EV biogenesis in fungi^25^. These include cell lysis processes reminiscent of those described in Gram-negative bacteria ^26, 64^. However, none of these mechanisms have been definitively validated or excluded ^26^. We reasoned that the collection of EV-production mutants generated in this study could be used to assess the contribution of cell death to EV biogenesis. To this end, we selected 45 mutants representing low, high, and WT EV producers and evaluated their cell viability during EV production on SD medium (Table S4). As shown in Fig. 5C, correlation analysis between cell viability and EV production revealed a slight but statistically significant negative correlation. However, whereas some poorly viable mutants, such as CNAG_00467, produced very high levels of EVs, some others with reduced viability, such as CNAG_04545, displayed EV production comparable to WT. Moreover, most EV-overproducing strains, as well as all low EV producers examined in this subset, exhibited no detectable defect in cell viability. Collectively, these observations indicate that cell viability alone is a poor predictor of EV production levels and suggest that cell lysis is unlikely to represent the sole mechanism underlying EV release in *C. neoformans*.

### Vesicles trafficking associated genes and genes implicated in stress responses are enriched in vesicles production regulators

GO-term enrichment analysis and manual functional annotation revealed enrichment of ESCRT components and multivesicular body (MVB) sorting pathway–associated GO terms among the negative regulators (Fig. 6A). Several mutants carrying deletions in genes coding for ESCRT^65^ proteins (Vps37, Vps28, Vps20, Vps22, Vps25, Vps36, Snf7 and Vps27) were among the strongest EV overproducers. Yet ESCRT mutant strains displayed no viability defect upon incubation at 30°C on SD for 24h (Table S4). Strikingly, some ESCRT proteins have also been previously shown to be implicated in EV production in *C. albicans* and/or *S. cerevisiae*, yet in these species they appear to act as positive regulators^15, 27^. We verified our screen results in *C. neoformans* by testing several independently prepared EV samples from individual ESCRT mutant strains (Table S2). We also re-introduced the WT gene in one ESCRT mutant and verified that this was associated with the restauration of a WT-level of both FLC susceptibility and EV production (Fig. S2). These apparently contrasting results with the ones previously obtained in *C. albicans* may reflect differences in growth conditions used for EV isolation in this later yeast (biofilm or liquid cultures), the strategies to evaluate EV concentration or distinct mechanisms governing EV production in these two evolutionary divergent yeasts. In fact, a similar negative regulation of EV production by ESCRT protein has been reported in mammalian HEK293T cells^66^.

**Figure 6.**
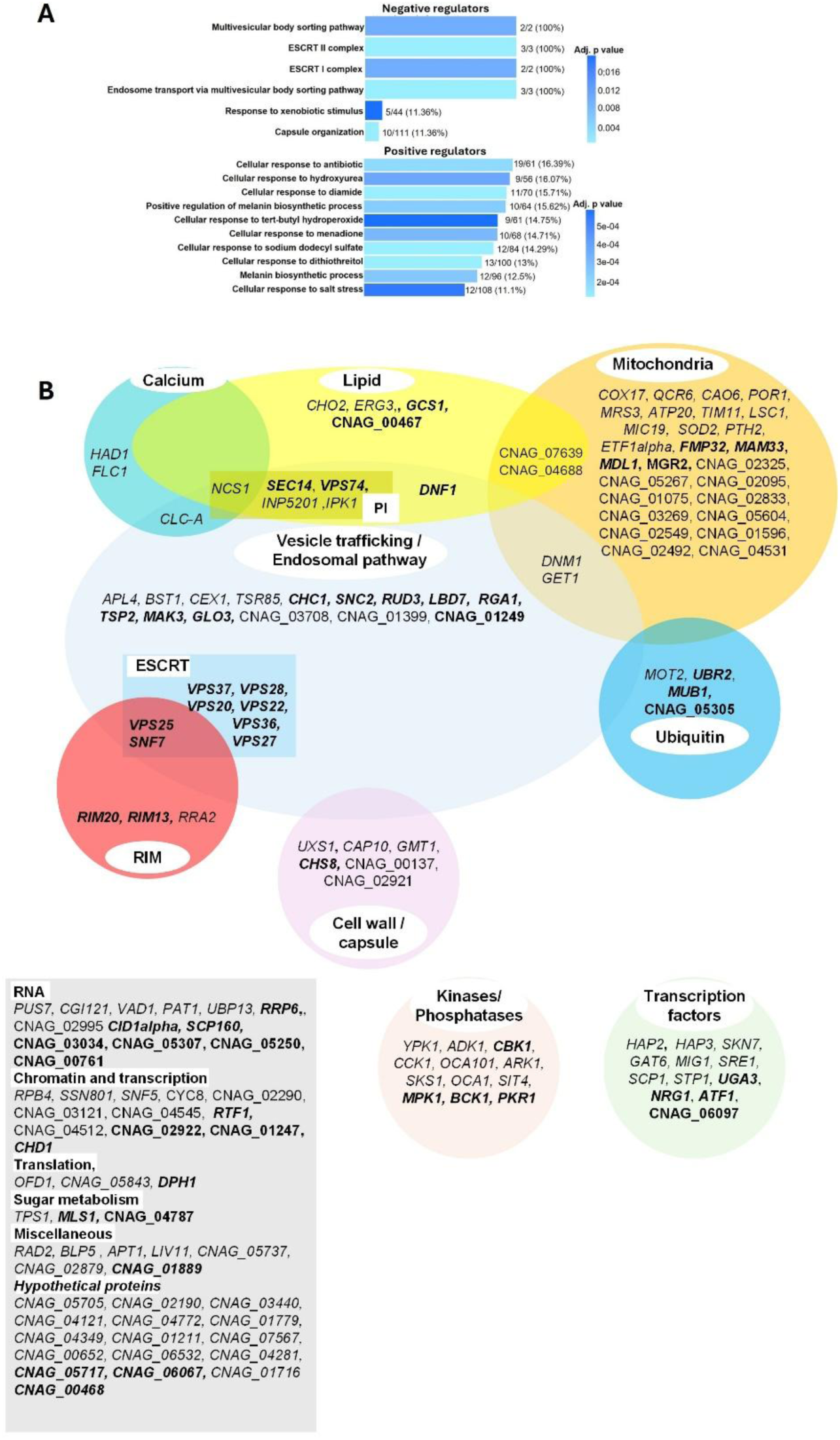
| The EV production regulators. **A.** Significantly enriched Gene Ontology (GO) terms among the EV production regulators, as identified using FungiFun3 with default parameters^42^. The percentage indicates the ratio of genes associated with a given GO term relative to the total number of background genes in that category. Negative regulators: GO:0071985: multivesicular body sorting pathway; GO:0000814: ESCRT II complex; GO:0000813: ESCRT I complex; GO:0032509: endosome transport via multivesicular body sorting pathway; GO:0042493: response to xenobiotic stimulus; GO:0045230: capsule organization. Positive Regulators: GO:0071236: cellular response to antibiotic; GO:0072711: cellular response to hydroxyurea; GO:0072738: cellular response to diamide; GO:0048023: positive regulation of melanin biosynthetic process; GO:0072736: cellular response to tert-butyl hydroperoxide; GO:0036245: cellular response to menadione; GO:0072707: cellular response to sodium dodecyl sulfate; GO:0072721: cellular response to dithiothreitol; GO:0042438: melanin biosynthetic process; GO:0071472: cellular response to salt stress. B. Functional annotation of the 158 genes regulating EV production revealed an enrichment for genes coding for vesicle trafficking and related pathways. Gene names correspond to the closest homologs in *S. cerevisiae*, unless a *C. neoformans* gene designation was already established. Genes were grouped into functional categories based on their predicted or experimentally validated functions. Of note, *GCS1* here refers to the ceramide glucosyltransferase encoding gene which is absent in *S. cerevisiae*. *APT1* refers to the adenine phosphoribosyltransferase encoding gene. The *C. neoformans APT1* encoding a flippase is here noted according to its reciprocal homolog in *S. cerevisiae DNF1*.

Consistent with a role of the ESCRT pathway in EV production regulation, several mutant strains affecting the Rim pathway, which mediates fungal response to alkaline pH and includes several ESCRT components also displayed altered EV production, being either positively (Rim13, Rim20) or negatively (Rra2) affected (Fig. 6B Table S2)^63, 67^. Interestingly, one also can note the presence of the three elements (Sre1, Scp1, Stp1) of the other alkaline pH response which is known to regulate the production and delivery of sterols to the plasma membrane to maintain appropriate cell homeostasis in fungi ^68^. This is also in good agreement with the presence of Erg11 and Erg3 in the EV regulon.

Twenty-four additional genes encoding proteins with predicted roles in intracellular vesicle trafficking and endosomal pathways emerged as regulators of EV production (Fig. 6B; Table S2), further supporting the central role of this pathway in EV biogenesis. Several of these proteins are associated with lipid biosynthesis. For example, we identified another flippase-encoding gene, here designated *DNF1* (also known as *APT1* in *C. neoformans*^48^) as a negative regulator of EV production in *C. neoformans*. We also found several genes involved in phosphoinositide (PI) metabolism (*SEC14, VPS74, INP5201, NCS1, IPK1*), which play critical roles in regulating membrane dynamics and vesicular transport^69^.

We also found in our screen 31 genes encoding proteins involved in mitochondrial metabolism (Fig. 6B; Table S2). Mitochondrial activity has also previously been linked to FLC susceptibility in pathogenic fungi^70, 71^ and we have previously shown that the Hap complex which is known to regulate mitochondria metabolism also regulates EV production in *C. neoformans*^16^. Hap2 and Hap5 were actually re-identified as strong positive regulators of EV production in the present analysis thus validating the quality of our screening. The identification of Mig1 which regulates mitochondrial function and FLC susceptibility in *C. neoformans*^70^ as a positive regulator of EV production is also in good agreement with central role that mitochondrial activity seem to play in regulating EV production.

As expected, several genes encoding cell-wall and capsule components, as well as genes associated with the ubiquitin pathway, emerged from the screen. The role of the fungal cell wall in regulating EV production has been discussed^26, 29^ but remains poorly characterized, while ubiquitin-related pathways are known to participate in endosomal trafficking and protein targeting to EVs in other systems^72, 73^.

Analysis of positive regulators of EV production identified several GO-terms associated with cellular responses to different stresses (Fig. 6A). Genes associated with these GO-terms are here mostly kinases and phosphatases implicated in diverse transduction pathways in fungi. Some of them are known direct regulators of the pathways identified above as involved in EV production regulation (Fig. 6B). Notably, the ceramide activated Sit4 serine-threonine phosphatase is known to control mitochondrial function, ER to Golgi traffic and cell wall organization in *S. cerevisiae*^74–76^. Similarly, the serine-threonine protein kinase; Ypk1 is required for plasma membrane lipid and protein homeostasis and regulate endocytosis^77^. We also confirmed the negative regulatory role of the serine-threonine protein kinases Bck1 and Mpk1 which encode components of the CWI pathway complex^29, 78^.

In agreement with the strong enrichment of genes associated with calcium homeostasis within the EV regulon and with observed impact of calcium depletion on EV production (Fig. 4), we identified three genes directly or indirectly implicated in calcium homeostasis among the EV regulators. Whereas deletion of *FLC1* in *C. neoformans* has been shown to be associated with elevated cytosolic calcium levels and increased nuclear localization of the calcineurin-dependent transcription factor Crz1^79^, the cryptococcal ortholog of neuronal calcium sensor 1 (*NCS1*) has been shown to regulate Ca^2+^ homeostasis under the control of the calcineurin signaling pathway^80^. Similarly, CLC-A is a reciprocal homolog of the *S. cerevisiae* Golgi voltage-gated chloride channel *CLC1* which has been shown to regulate calcium homeostasis in *Cryptococcus deneoformans*^81^.

Overall, although only six genes (i.e. *BCK1*, *SRE1*, the chitin synthase-encoding gene *CHS8*, the tetraspanin-encoding gene *TSP2*, *ERG3* and CNAG_02921, which encodes a GPI-anchored protein of unknown function) are shared between the EV regulon and the EV production regulator gene sets, the pathways represented in the two sets show striking similarities (Fig. 3 and Fig. 6B). In particular, both sets contain numerous genes involved in vesicular trafficking, consistent with the established role of multivesicular bodies and the endosomal pathway in EV biogenesis in metazoans^82^. The substantial representation of genes involved in lipid, cell wall and mitochondrial metabolism in both sets further implicates these processes in EV biosynthesis. Together, these findings identify vesicular trafficking as a central regulatory axis of EV production while highlighting additional contributions from lipid, cell wall and mitochondrial metabolism. Finally, consistent with the composition of the EV regulon, we identified key regulators of calcium homeostasis among the major regulators of EV production, further implicating calcium signalling in the control of EV biogenesis.

### *HAD1* is a positive regulator of EV biogenesis in *C. neoformans*

In 2016, Park and colleagues used phosphoproteomic and quantitative mass spectrometry approaches to identify 44 proteins as substrates of calcineurin in *C. neoformans*^83^. Two of these proteins were among the EV regulators identified in our study. Snf7 is a negative regulator of EV production and a component of the ESCRT complex^65^. Its presence confirmed the pre-established link between vesicles trafficking and calcineurin pathway. The second is Had1 and it caught more specifically our interest for several reasons. First, its interaction with Cna1 has been studied in detail^84, 85^, further supporting its connection to the calcineurin pathway. Second, Had1 was previously shown to contribute to thermotolerance, cell wall integrity and virulence in *C. neoformans*^84^. Third, the *had1Δ* mutant strain was one of lowest EV producer (Table S2).

We confirmed the EV production phenotype of *had1Δ* using both a rapid EV isolation protocol and the conventional protocol widely used in the fungal field, which includes an ultracentrifugation step^62^. The *had1Δ* strain produced approximately 50-fold fewer vesicles than both the WT and reconstructed (*had1Δ* + *HAD1*) strains (Fig. 7A). We also confirmed that *had1Δ* was more resistant to FLC than the WT and reconstructed strains, with MIC_50_ values of 12.6 and 8.4 μg/mL, respectively (Fig. 7B). We noticed that *HAD1* deletion resulted in a modest growth defect on SD medium after 24 h at 30 °C (Fig. S3). However, extending the growth period to 28 or 30 h did not restore EV production in the *had1Δ* mutant to WT levels, supporting a major role for *HAD1* in the regulation of EV production in *C. neoformans* (Fig. S3). We also observed that cells grown under EV isolation conditions were smaller in the *had1Δ* strain (median diameter 3.95 μm) than in the WT (5.1 μm) and reconstructed (5.06 μm) strains (Fig. 7C,D). Interestingly, the *had1Δ* strain also produced EVs with a modest but consistent reduction in diameter compared with the WT and reconstructed strains (Fig. 7E). Together, these findings indicate that *HAD1* influences not only the amount of EVs produced but also their size, suggesting a broader role in the regulation of EV biogenesis.

**Figure 7.**
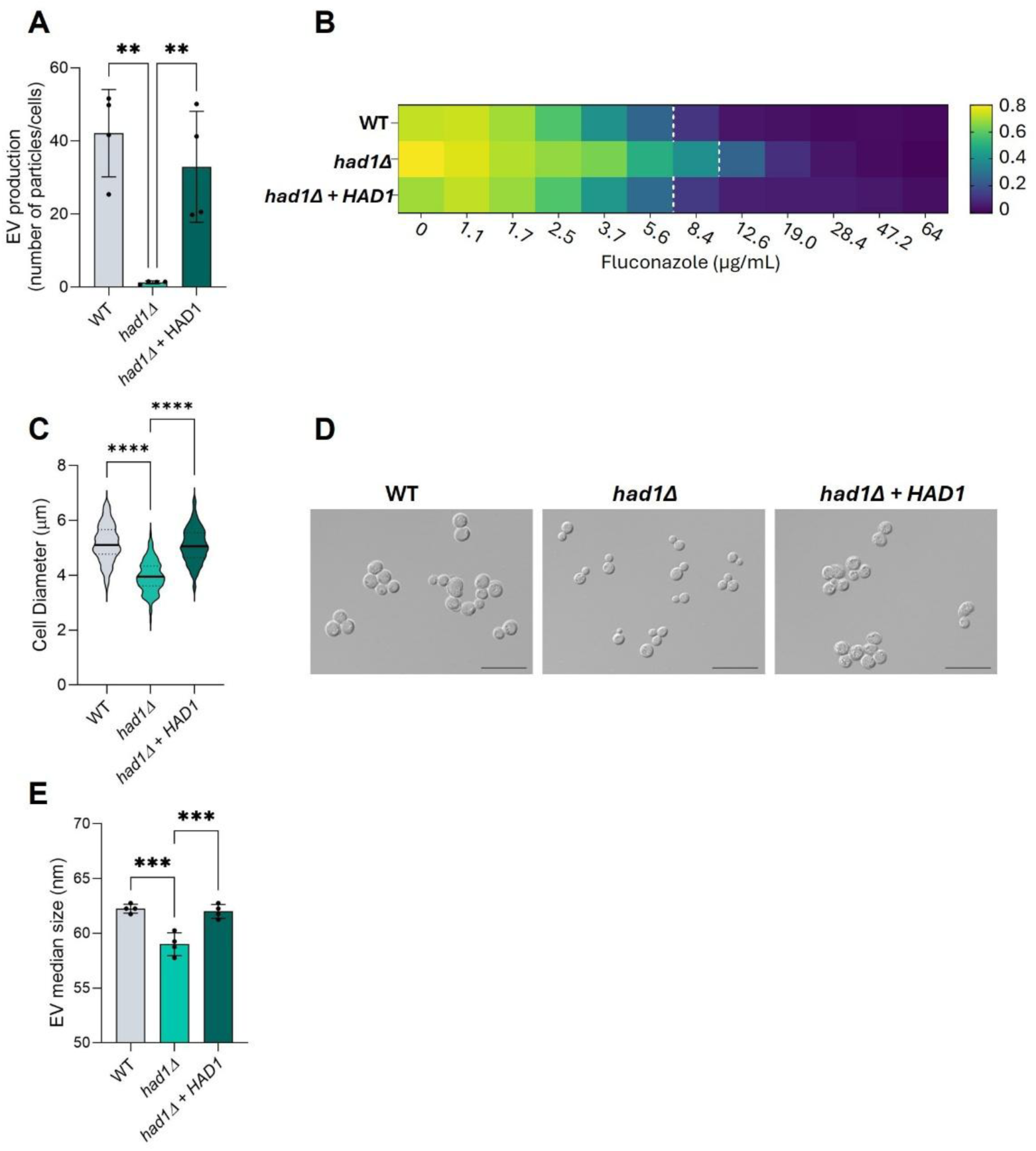
| Had1 regulates EV production in *C. neoformans*. **A.** Quantification of EVs released by WT, *had1Δ*, and *had1Δ* + *HAD1* strains using the ultracentrifugation protocol, measured by nanoFCM. Data represent means ± SD \*\**p* < 0.01 (ANOVA followed by Tukey’s multiple comparisons test). **B.** Heat map of the FLC MIC assay. The white line indicates the MIC_50_ (n = 6). **C.** Cell size differences associated with the *had1Δ* mutation. Cells were grown under EV-producing conditions on SD agar plates for 24 h at 30 °C and imaged using an Olympus IX83 microscope. Cell diameters were quantified using ImageJ (n = 150). Error bars show means ± SD.**** *p* <0.0001 (ANOVA followed by Tukey’s multiple comparisons test). **D.** Representative images of each strain. Scale bar correspond to 20µm. **E.** Median EV diameters measured by NanoFCM. Error bars show medians ± SD.*** *p* <0.001 (ANOVA followed by Tukey’s multiple comparisons test).

### Calcineurin pathway regulates EV biogenesis in *C. neoformans*

Our data suggested that the calcineurin pathway regulates EV production in *C. neoformans*. To verify this hypothesis, we recovered the knockout mutant strains for both calcineurin subunits (i.e. the catalytic subunit A Cna1 and regulatory subunit Cnb1) from the Madhani collection, validated them by WGS and tested their EV production. As expected, for both mutant strains, we observed a strong reduction in EV release (Fig. 8A). Because both mutant strains displayed a reduced growth on SD medium (Fig. S4), we extended the time of growth to 28 and 30 hours, but we did not observe noticeable modification of the EV production phenotype (Fig. S4). We also observed a strong reduction of EV production in the presence of sub-inhibitory concentration of two calcineurin pharmacological inhibitors, cyclosporin A and FK506 confirming the regulatory role of calcineurin pathway in EV production in *C. neoformans* (Fig 8B,C).

**Figure 8.**
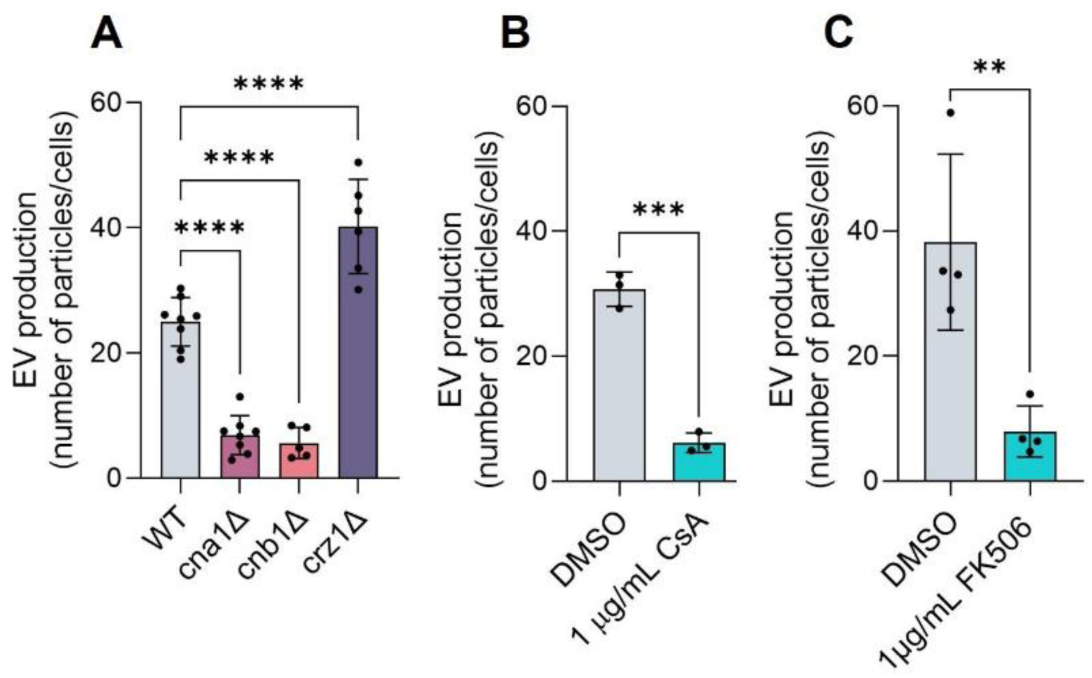
| The calcineurin pathway regulates extracellular vesicle (EV) production. **A.** Quantification of EV production by WT, *cna1Δ*, *cnb1Δ* and *crz1Δ* mutants using nanoFCM. (**** *p* < 0.0001, one-way ANOVA followed by Dunnett’s multiple-comparisons test). **B.** Quantification of EV production by WT cells treated with the calcineurin inhibitor cyclosporin A (CsA) using nanoFCM. (** *p* < 0.01; *** *p* < 0.001, unpaired two-tailed Student’s *t*-test). **C.** Quantification of EV production by WT cells treated with the calcineurin inhibitor FK506 using nanoFCM. (** *P* < 0.01; *** *p* < 0.001, unpaired two-tailed Student’s *t*-test).

The most studied calcineurin target in fungi is the transcription factor Crz1 which shuttle from the cytoplasm to the nucleus upon Cna1-dependant dephosphorylation^57^. Strikingly, the *crz1Δ* mutant strain recovered from Madhani collection and validated through PCR-assays produced more EV than the WT strains after 24, 28 and 30h suggesting a negative regulatory role of this transcription factor (Fig. 8A). This result also suggests that Crz1 and Had1 do not act in the same pathway to regulate EV production.

### *HAD1* encodes is a glycerol-3-phosphate phosphatase

Had1 is a member of a group of four closely related *C. neoformans* proteins (Had1, Had6 [CNAG_06122], Had8 [CNAG_06132] and Had9 [CNAG_06698]), which are all annotated as phosphatases in FungiDB. BLAST analysis identified the closest *S. cerevisiae* homolog of Had1 as Dog1, a 2-deoxyglucose-6-phosphate phosphatase, with which it shares 18.01% amino-acid identity. Dog1 also represents the best *S. cerevisiae* match for Had6, whereas Had8 and Had9 exhibit higher similarity to the glycerol-3-phosphate phosphatase Gpp1. However, none of these BLAST hits is reciprocal, and the biological functions of the Had proteins in *C. neoformans* have remained unresolved.

To address this question, we first performed a three-dimensional similarity search using VAST^86^, which identified *S. cerevisiae* Gpp1 as the top hit (32.1% identity). The predicted structures of Had1 and Gpp1 are indeed highly similar, although the AlphaFold3^87^ predictive model of Had1 contains a long N-terminal intrinsically disordered region that is absent from Gpp1 (Fig. 9a). This structural correspondence is consistent with recent studies proposing that Had1 functions as a glycerol-3-phosphate phosphatase, based on phenotypic evidence including pronounced sensitivity to NaCl^88–90^ reminiscent of the phenotype of a glycerol-3-phosphate phosphatase-deficient *S. cerevisiae* mutant^91^. To test this hypothesis directly, we generated a conditional *S. cerevisiae* double mutant (tetOFF-*gpp1 gpp2Δ*), transformed with a centromeric plasmid expressing *HAD1* under the control of the *S. cerevisiae TDH3* promoter, and assessed growth on YPD medium supplemented with 500 mM NaCl in the presence or absence of doxycycline. As presented on Fig.9B, expression of Had1 fully restored NaCl resistance in the *S. cerevisiae* glycerol-3-phosphate phosphatase-deficient strain. Moreover, HPLC-based quantification intracellular glycerol-3-phosphate concentration in *C. neoformans* revealed that *had1Δ* cells accumulated approximately threefold higher levels of glycerol-3-phosphate than the WT cells (Fig. 9C), further supporting the conclusion that Had1 functions as a glycerol-3-phosphate phosphatase.

**Figure 9.**
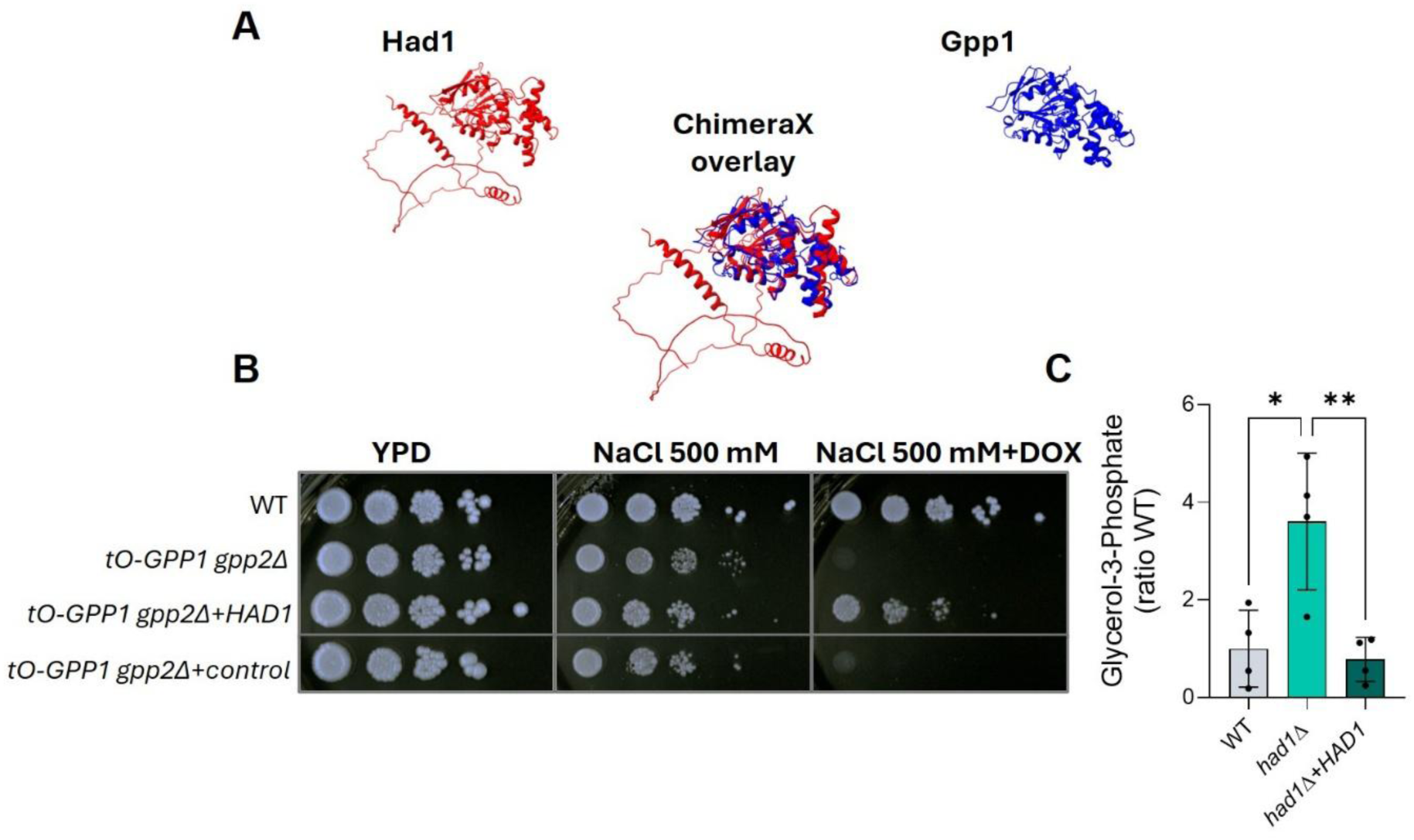
| Had1 is a glycerol-3-phosphate phosphatase. **A.** Structural comparison of Had1 and the *S. cerevisiae* glycerol-3-phosphate phosphatase Gpp1, as predicted by AlphaFold3. Overlay was generated using UCSF ChimeraX^92^. **B.** Spot assays of *S. cerevisiae* WT and *tetOFF-GPP1 gpp2Δ* strains expressing *C. neoformans HAD1* or carrying an empty vector, grown in the presence or absence of doxycycline (DOX). Pictures were taken after 3 days of incubation at 30°C. **C.** Quantification of intracellular glycerol-3-phosphate in WT, *had1Δ* and *had1Δ* + *HAD1* strains by HPLC. Error bars indicate mean ± SD. (*p* < 0.05; *p* < 0.01 (ANOVA followed by Tukey’s multiple-comparisons test).

### Had1 plays a unique role in the early steps of the phospholipid synthesis pathway to regulate EV production

In *S. cerevisiae*, the first step in phospholipid biosynthesis is the acylation of glycerol-3-phosphate (G3P) to lysophosphatidic acid (LPA). LPA is then acylated to form phosphatidic acid (PA) by the 1-acyl-sn-G3P acyltransferases Slc1p and Ale1p. G3P acyltransferases (Sct1p and Gpt2p) catalyze the initial step in both branches of the pathway by acylating G3P to form either LPA or 1-acyl-DHAP from dihydroxyacetone phosphate (DHAP). Sct1p and Gpt2p also catalyze DHAP acylation to generate 1-acyl-DHAP, which is then reduced to LPA by the 1-acyl-DHAP reductase Ayr1^93, 94^ (Fig. 10A). We retrieved the corresponding *C. neoformans* mutant strains and confirmed their identity by whole-genome sequencing. It should be noted that the *SCT1*/*GPT2* homologue CNAG_01711 was absent from the Madhani deletion collection and that repeated attempts to delete this locus were unsuccessful. This suggests that it is essential, which is consistent with previous saturation transposon mutagenesis analyses^95^. We next used NanoFCM to assess EV production in these mutants, as well as in deletion mutant strains of the other *HAD* genes. Strikingly, none of the strains exhibited a significant defect in EV production, suggesting a unique role for Had1 in regulating this process (Fig. 10B).

**Figure 10.**
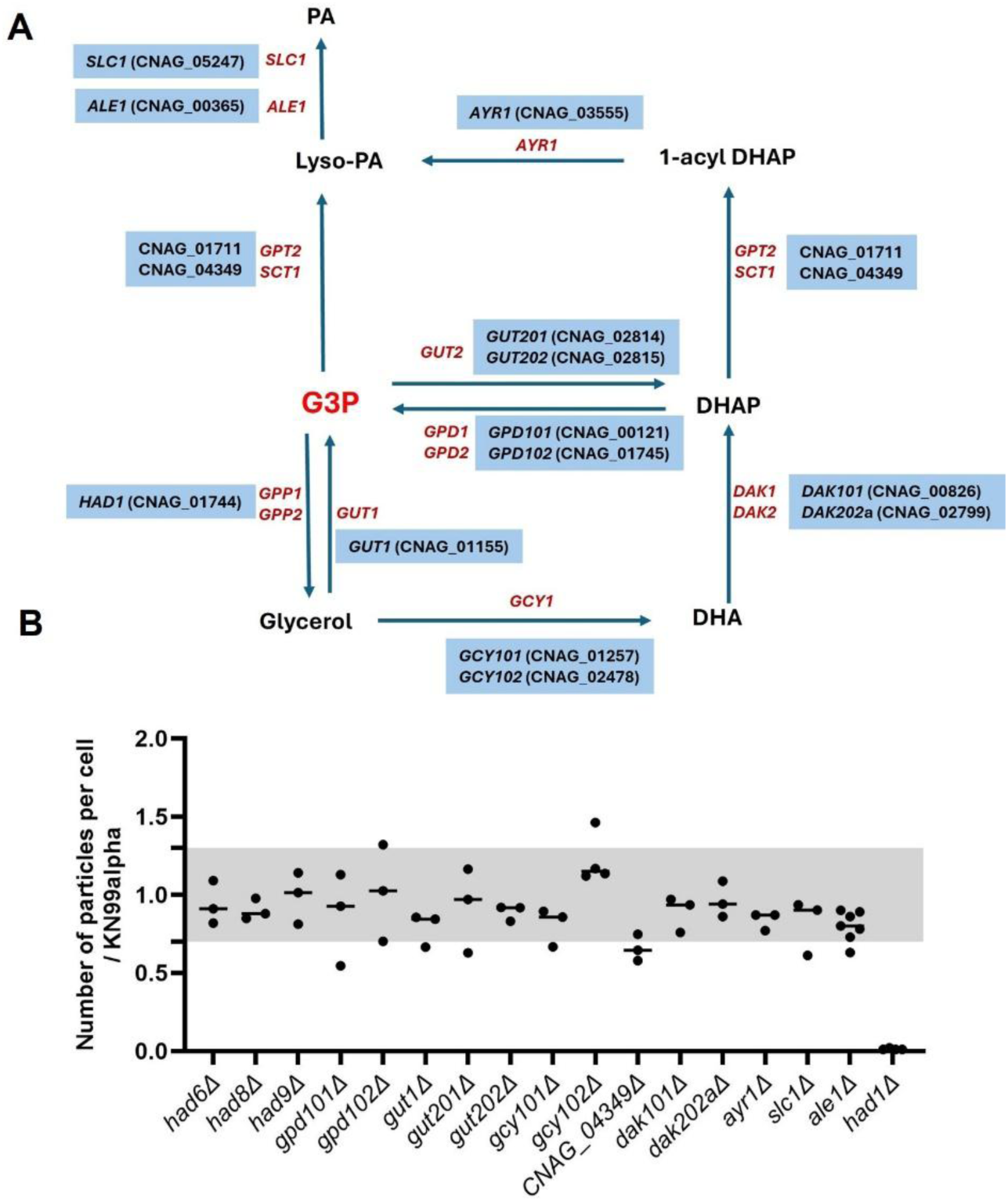
| Impact of the deletion of genes implicated in glycerol 3 phosphate (G3P) metabolism on EV production. **A.** Schematic of genes and metabolites involved in G3P metabolism as characterized in *S. cerevisiae*. *S. cerevisiae* gene names are shown in brown, and homologs in *C. neoformans* are highlighted in blue boxes. The positions of glycerol, glycerol-3-phosphate (G3P) lysophosphatidic acid (Lyso-PA), phosphatidic acid (PA), dihydroxyacetone phosphate (DHAP), dihydroxyacetone (DHA) and a 1-acyl-*sn*-glycerol 3-phosphate (1-acyl-DHAP) in this pathway are indicated. **B.** Quantification by NanoFCM of EV release in knockout mutants of genes involved in G3P metabolism.

### Lack of the glycerol-3-phosphate phosphatase reshape the cellular lipid profile

Glycerol-3-phosphate phosphatases catalyze the conversion of glycerol-3-phosphate to glycerol. Theoretically, *HAD*1 deletion should reduce the intracellular concentration of glycerol. However, supplementing the growth medium with glycerol did not suppress the FLC-resistant phenotype or reduced vesicle production of the *had1Δ* strain (see Fig. S5), suggesting that these traits are unlikely to be caused by a simple reduction in intracellular glycerol levels. *HAD1* deletion also leads to the accumulation of glycerol-3-phosphate, an early intermediate in phospholipid biosynthesis. This suggests that loss of *HAD1* may disrupt lipid homeostasis in *C. neoformans*. To test this hypothesis, we performed lipidomic profiling using whole lipid cell extracts from the WT and *had1Δ* mutant strains. Analysis of the data supported our hypothesis, revealing a striking alteration in the cellular lipid composition of the *had1Δ* mutant, characterized by an accumulation of structural lipids (Fig. 11, Table S5). Specifically, we observed significant upregulation of several major phosphatidylcholine (PC) and phosphatidylethanolamine (PE) species, indicating a substantial increase in the structural phospholipid (PL) pool. While levels of the upstream metabolite lysophosphatidic acid (LPA) remained unchanged, this PL surplus was accompanied by a striking elevation of multiple downstream lysophospholipid classes, including Lyso-PC, Lyso-PE, Lyso-PG, Lyso-PI and Lyso-PS, as well as a significant increase in monoglycerides (MGs). This extensive lipid cleavage concurrently led to a broad upregulation of free fatty acids (FFAs) and the subsequent accumulation of short- and medium-chain acylcarnitines. Together, these lipid changes suggest that an initial overproduction of structural PLs in the *had1Δ* triggers massive compensatory lipolytic degradation and severe biophysical alteration of membrane packing, which correlates with a strong reduction in EV production.

**Figure 11.**
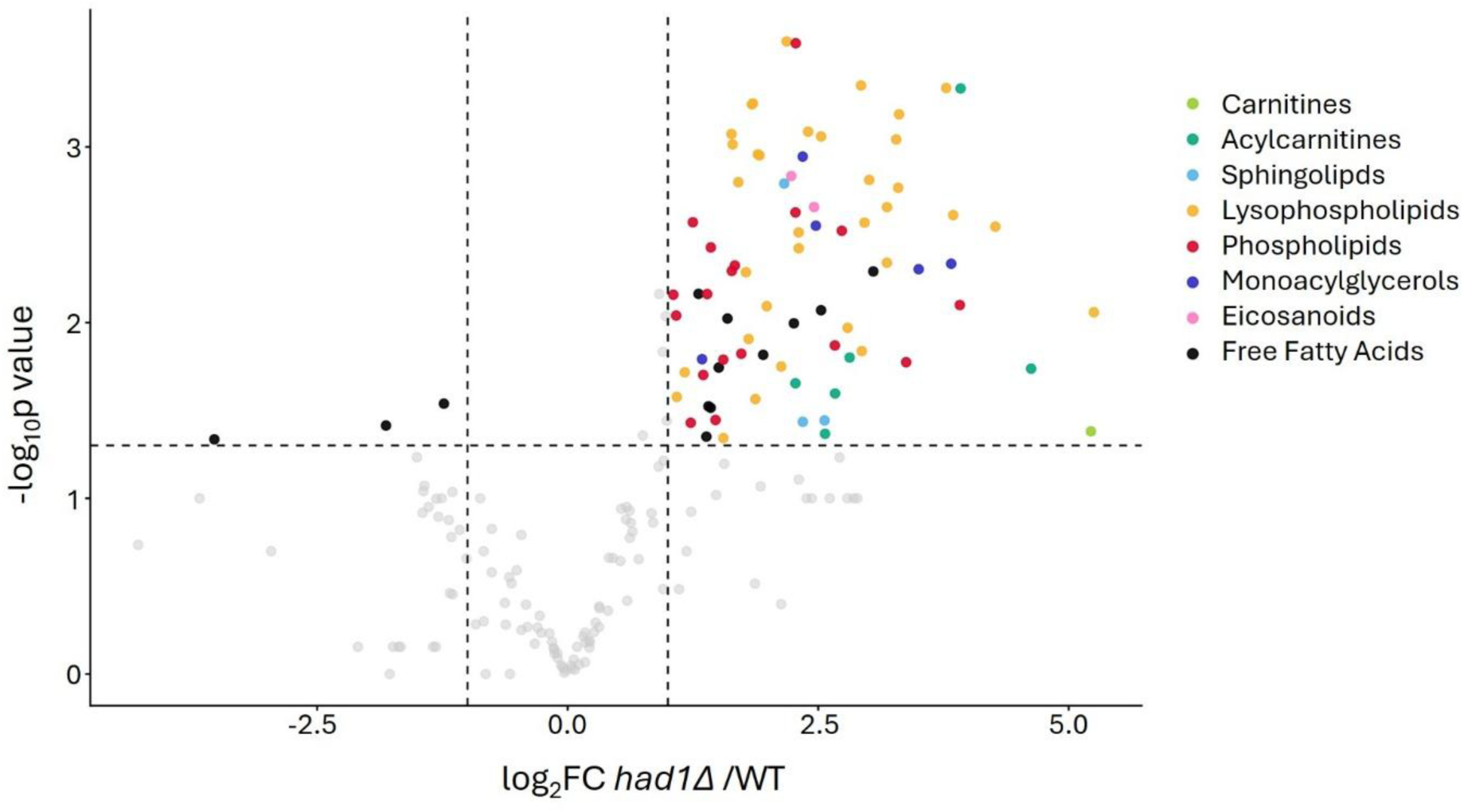
| *HAD1* deletion is associated with altered lipid metabolism. Volcano plot comparing lipidomic profiles of *had1Δ* and WT cells. The x-axis shows log₂ fold changes in lipid abundance between strains, and the y-axis indicates significance (–log₁₀ *p*-value). Each point represents a detected lipid species.

## Discussion

### A first global view of EV regulatory pathways in fungi

Whereas interest in microbial, and more specifically fungal, EVs has grown substantially in recent years, our understanding of their biosynthetic pathways and regulatory mechanisms remains limited. To date, only a handful of genes have been convincingly shown to regulate either the quantity or the composition of fungal EVs^15, 16, 27^ ^29, 96, 97^. Here, we provide the first comprehensive analysis of EV regulators and associated regulon genes in fungi with implications across microbial systems.

Consistent with observations in mammalian systems^25, 82, 98^, analysis of these gene sets supports a central role for endosomal pathway–related genes beyond the canonical ESCRT machinery in EV production in fungi providing further support to previous hypotheses^26^. In addition, our data reveals a prominent role for mitochondrial metabolism in the regulation of EV production. Our previous identification of the Hap complex as a major regulator of EV production together with the growth phase specific production of EVs already led us to suggesting a central role of this organelle in EV production regulation^16^. A similar association has been reported in mammals, where mitochondrial metabolism has also been identified as a regulator of EV secretion in several genome-wide screens performed in different human cell lines^66, 99^. The recovery of regulatory pathways previously identified in independent mammalian screens provides cross-system validation of our screen, supporting the robustness and biological relevance of the identified EV regulators.

In mammals, multiple mechanisms have been proposed to account for mitochondria biology in EV production regulation, including TRX2-dependent ubiquitination and degradation of the small GTPase Rab35, which impairs exosome secretion^100^, as well as alterations in lipid metabolism mediated by mitochondria-associated endoplasmic reticulum membranes^101, 102^. In fungi, although the mitochondrial metabolism is already known to regulate azole resistance through multiple mechanisms^103^, how it influences EV production remains unclear. In light of our findings, one possibility is that this regulation involves the established role of mitochondria as dynamic Ca²⁺ reservoirs and regulators of intracellular calcium homeostasis^104^. Notably, mitochondrial dysfunction modulates azole susceptibility through a calcium signalling-dependent pathway in *A. fumigatus*^105^. The identification of calcineurin-dependent regulation of EV production in our study raises the possibility of a functional link between mitochondrial metabolism, calcium signalling, azole susceptibility and EV production in *Cryptococcus*.

We also identified multiple capsule- and cell wall–associated genes as regulators of EV production. A role for capsule structure in fungal EV regulation has been largely discussed previously^27, 29, 106–108^. In agreement with this hypothesis, several cell wall mutants, including the chitin synthase mutant *chs3Δ*, have been shown to produce increased levels of EVs compared with the WT strain in *S. cerevisiae*^27^. Similarly, in *Mycobacterium tuberculosis*, alterations in cell wall integrity modulate EV production, suggesting that cell wall–dependent regulation of EV biogenesis may represent a conserved mechanism in other cell-walled organisms^108^.

### A model for calcineurin-dependent regulation of EV production

Our transcriptomic and genetic analyses further implicate the calcium–calcineurin signalling pathway in the regulation of EV production in *Cryptococcus*. Calcium influx has been widely reported to regulate the biogenesis and release of both exosomes and microvesicles in mammalian systems^109–111^. This process has been proposed to involve phospholipid scramblases, calcium-activated proteases (calpains) and components of the ESCRT machinery^11^. Notably, all of these factors were identified in our screen, providing further support for the involvement of calcium-dependent mechanisms in EV production in fungi.

We propose a model in which changes in extracellular calcium availability, or alterations in intracellular calcium concentrations arising through diverse mechanisms, including changes in mitochondrial metabolism^104^ regulate the phosphorylation state of the glycerol-3-phosphate phosphatase Had1 through calcineurin signaling (Fig. 12). Our data indicate that this regulation is independent of the transcription factor Crz1, consistent with previous observations of Crz1-independent calcineurin responses during thermotolerance and mating^57, 112, 113^. Interestingly, Crz1 appears to have an opposing effect on EV production, suggesting that calcineurin-dependent regulation of EV biogenesis may be subject to feedback control.

**Fig. 12.**
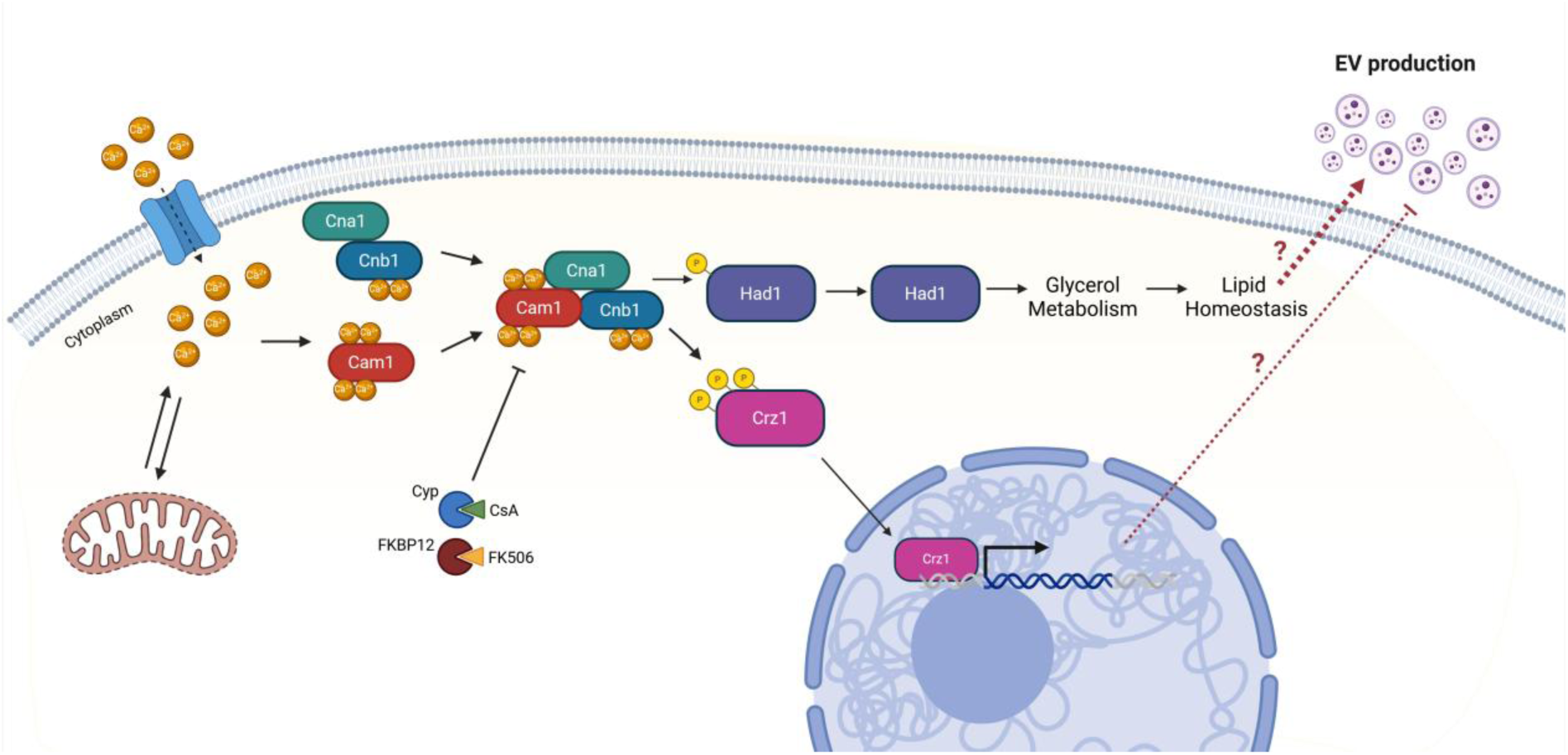
Model of Ca^2+^/calcineurin influence in EV biogenesis. Ca^2+^/calcineurin pathway acts as a modulator of EV production, with one target, Had1, as a positive regulator and the second target, Crz1, as a negative regulator.

Changes in Had1 phosphorylation could, in turn, alter its subcellular localization and/or enzymatic activity, thereby modulating intracellular glycerol-3-phosphate (G3P) levels. Strikingly, *had1Δ* was the only mutant of the early phospholipid metabolic pathway that displayed a pronounced EV production phenotype. Although, we cannot fully exclude the possibility that this striking phenotype reflects limited functional redundancy for *HAD1*, our lipidomic analyses suggest that the accumulation of the G3P is indeed responsible for this phenotype. In this model, G3P accumulation overloads the Kennedy pathway^94^, leading to overproduction of membrane phospholipids (PE and PC) and alteration of membrane structure and composition. While the mechanisms linking these alterations in lipid composition to EV production remain to be established, it is known that perturbations in lipid homeostasis impair EV production in mammalian systems^25, 114^. Further support for a connection between lipid metabolism and EV production comes from the strong co-regulation observed in our dataset between susceptibility to FLC and EV production in *C. neoformans*.

### Limitations of the approach

Our mutant screening approach has several limitations and may have led to both false-positive and false-negative identifications. First, although the pre-screening strategy proved highly effective, as demonstrated by the strong enrichment of mutants impaired in EV production among FLC-resistant or -sensitive strains, it was not exhaustive. We identified multiple mutants with altered FLC susceptibility that did not exhibit detectable EV production defects. Conversely, we found that calcineurin pathway mutants displayed impaired EV production but were not identified as FLC-resistant or -sensitive in our pre-screen. This discrepancy may stem from the strategy used to identify FLC susceptibility mutants. Indeed, *cna1Δ* and *cnb1Δ* have been identified as resistant using competitive growth assays at low FLC concentrations by the Madhani group^63^. Second, the extensive phenotypic characterization of the Madhani mutant collection suggests that some observed phenotypes may not be directly attributable to disruption of the targeted gene but instead to secondary effects on downstream or upstream gene regulation. We identified nine such gene pairs among our EV production–mutant strains, for which the bona fide EV regulator remains unclear. Third, although the Madhani collection encompasses a large fraction of non-essential genes, it is not complete. Several genes known to regulate FLC susceptibility, and whose functions suggest potential roles in EV biogenesis or regulation, such as *CDC50*, *PDR16*, *ARF1*, and *CFT1*, are absent from this collection. Actually, *CDC50* has been recently identified as a regulator of EV production in *Cryptococcus*^49^. These observations indicate that our identification of EV regulatory genes is likely incomplete and underscore the need for additional, complementary approaches to fully elucidate the genetic network governing EV production in *C. neoformans*.

## Conclusion and perspectives

This study establishes a foundation for understanding EV biosynthetic pathways in fungi and highlights the power of combining forward genetic screening with time-course RNA-seq analyses. We propose that changes in mitochondrial metabolism during the transition from exponential to stationary phase alter intracellular calcium levels, which are transduced through the calcineurin pathway, leading to changes in glycerol-3-phosphate abundance and lipid homeostasis, ultimately modulating EV production. However, within this model, the mechanistic link between altered lipid homeostasis and EV biogenesis remains unresolved. In particular, it is unclear which modifications of plasma or intracellular membranes are critical and which molecular mechanisms mediate these effects. Moreover, the specific EV biogenesis pathway regulated under these conditions remains unknown. By analogy with mammalian systems, the existence of exosome-like and microvesicle-like pathways has been proposed in fungi^26^ and multiple microscopy- and genetics-based studies support this hypothesis^15, 96, 97, 115–117^. The remarkable diversity in EV size, morphology, and surface features observed by cryo-electron microscopy further suggests the involvement of distinct EV production routes. In addition, we recently reported the generation of apoptotic-like vesicles in CWI mutant strains^29^. For example, the reduced size of EVs produced by the *had1Δ* mutant strain may result from altered lipid composition and/or the preferential production of a specific EV subtype. Future studies will be required to delineate the regulation and functional roles of these distinct EV biosynthetic pathways and to define their contributions to biology and virulence of pathogenic fungi.

## Materials and methods

### Media and culture conditions

All the strains used in this study are from G. Janbon laboratory collection and are stored at -80 °C in glycerol 40%. Before each experiment, the strains were plated in YPD-agar plates (1% yeast extract, 2% bacto-peptone, 2% dextrose, and 2% bacto-agar) and incubated at 30°C for 48 h. Cell inoculum was performed in YPD broth (1% yeast extract, 2% bacto-peptone, 2% dextrose) and to isolate EVs, cells were plated in SD-agar plates (0.67% yeast nitrogen base without amino acids (Difco), 2% glucose (Sigma-Aldrich) and 2% bacto-agar (BD Biosciences)). In some experiments, SD or YPD-agar plates were supplemented with Cyclosporine A (CsA; 30024, Sigma–Aldrich), FK506 monohydrate (F4679, Sigma–Aldrich), BAPTA tetrapotassium salt (1,2-bis(o-aminophenoxy)ethane-N,N,N′,N′-tetraacetic acid, B1204, Invitrogen) or EGTA (ethylene glycol-bis(β-aminoethyl ether)-N,N,N′-tetraacetic acid, E3889, Sigma–Aldrich). For strains unable to grow in SD-medium, cells were plated in SC-agar plates (0.67% yeast nitrogen base without amino acids (Difco), 0.2% complete supplement mixture (Formedium), 2% glucose (Sigma-Aldrich) and 2% bacto-agar (BD Biosciences)). The concentration of each compound is indicated in each figure legend.

### Fungal strains and mutant construction

The knockout library has been constructed by Hiten Madhani lab (UCSF, USA) and obtained from the Fungal Genetic Stock Center. The genotypes of the mutant strains explored in this paper were confirmed by PCR using gene-specific internal primers and/or WGS. The *HAD1* gene were re-introduced into *MATα had1Δ::NAT* strain using a transient CRISPR-Cas9 expression system^118^. DNA fragments covering a genomic region from 600 bp upstream to 600bp downstream the ORF of the gene CNAG_01744 were PCR amplified and cloned into the pSDMA57 plasmid before being integrated at the genomic “safe haven” locus in *had1Δ*^119^. All plasmids and primer sequences used are provided in Tables S6 and S7, respectively.

For the *S. cerevisiae* complementation assay, a tetOFF-*gpp1 gpp2Δ* conditional expression double mutant was constructed. For that, the BY4741 *MAT***a** *gpp2Δ* strain from the systematic deletion collection was used^120^. The KanMX resistance cassette was replaced by an hygromycin resistance cassette using pAG32^121^. Then, the G418 resistance cassette and tTA-*ADH1*t-tetO2 module was amplified from pCM224 vector^122^ using primers designed with specific flanking sequences targeting upstream the start position of *GPP1* (YIL053W) coding sequence. The *S. cerevisiae gpp2Δ* was transformed and individual clones were tested by PCR on genomic DNA to test the correct insertion of the cassette. To construct the plasmid expressing *C. neoformans HAD1*, the specific cDNA was obtained from *C. neoformans* KN99α total RNA using Superscript IV (Invitrogen), following the manufacturer instructions.

The *HAD1* coding sequence was cloned in single copy CEN plasmid (*LEU2*) between a *TDH3* promoter and *ADH1* terminator. *S. cerevisiae* tetOFF-*gpp1 gpp2Δ* was transformed with this plasmid and positive transformants were selected on synthetic complete medium lacking leucine. For phenotypic assays, serial dilutions of cells were spotted on YPD medium and YPD containing NaCl (500mM) and/or doxycycline (10 µg/mL). Plates were incubated at 30°C for 72h and imaged using Phenobooth (Singer Instruments, Watchet, UK).

All strains studied are described in Table S8.

### FLC susceptibility tests

#### Screening spot assay

Cells from glycerol stocks (20 µL) were inoculated into 1 mL of liquid YPD in deep-well plates and grown for 48 h at 30°C with agitation. Cultures were then centrifuged, and cell pellets were washed and resuspended in 1 mL of sterile H₂O. Aliquots of 100 µL were transferred to 96-well plates and spotted onto YPD agar plates (growth control) and YPD agar plates supplemented with fluconazole (FLC) at 8, 16, or 32 µg/mL using a ROTOR+ robot (Singer Instruments, Watchet, UK). Plates were incubated at 30°C and imaged after 1, 2, and 5 days using a PhenoBooth (Singer Instruments, Watchet, UK). FLC-resistant and FLC-sensitive candidates were identified by visual inspection. Candidates identified in the initial screen were subjected to a second screening using the same experimental procedure to confirm their phenotypes.

#### Minimum Inhibitory Concentration (MIC)

MIC assays were performed as previously described^123^ with modifications. Briefly, cells from YPD plates were inoculated into 1 mL of liquid YPD in deep-well plates and grown for 24 h at 30°C with shaking. Cells were washed twice with sterile water, and approximately 2 × 10⁵ cells were inoculated into 96-well plates containing 125 µL of YPD supplemented with FLC at concentrations ranging from 1.1 to 64 µg/mL in a 1.5-fold dilution series. Untreated wells (0 µg/mL FLC) were used as controls. Plates were incubated at 30°C with shaking at 200 rpm. Cell density was measured after 48 h using a Tecan microplate reader (Tecan Trading AG, Switzerland). The MIC₅₀ was defined as the lowest FLC concentration resulting in 50% growth inhibition relative to the untreated control.

FLC disk diffusion assays were performed as previously described^16^.

### Time course RNA-Seq

#### RNA sample collection and library preparation

Cells were grown under EV-producing conditions on SD agar plates for 14, 16, 18, 20, or 24 h at 30°C. Cells were gently collected using a sterile loop and resuspended in cold water. Cell suspensions were then centrifuged at 6,000 rpm for 5 min, and the resulting pellets were immediately snap-frozen in liquid nitrogen. Total RNA was extracted as previously described^124^ RNA-Seq libraries were prepared from 1.5 µg of total RNA by polyA selection using the TruSeq Stranded mRNA kit (Illumina) according to the manufacturer’s instructions. Then, 100 bases were sequenced from both ends using an Illumina NextSeq500 instrument according to the manufacturer’s instructions (Illumina). Each condition was analyzed using three biological replicates.

#### Sequencing read trimming and mapping

Paired-end sequencing reads were trimmed, Illumina TruSeq adapter sequences were removed, and rRNA sequences were filtered out as previously described^16^. Cleaned reads were aligned to the H99 reference genome (NCBI Genome Assembly GCA_000149245.3) with Tophat2 v2.0.14^125^ using previously described parameters^16^.

#### Time course RNA-seq analysis

The differentially expressed genes were assessed using the DEseq2 v1.16^126^. Growth phase-regulated genes (fold change ≥ 2 and R-squared ≥ 0.75) were then identified based on the generalized additive model, and their normalized counts across different time courses were used to group genes according to the DP_GP_cluster method ^127^ using default parameters as previously described in https://github.com/PrincetonUniversity/DP_GP_cluster/blob/master/README.md.

### EV isolation and quantification

EVs were isolated from cells grown on solid SD media as previously described ^62^. Depending on the experiment, EV concentrations were determined by NanoAnalyzer (nanoFCM) equipment following manufacturer protocols. TE filtered at 0.02-μm was used to define the event triggering threshold. Samples were diluted in 0.02-μm-filtered TE.

### Screening strategy

Each FLC-resistant or FLC-sensitive strain identified by spot assay was first assessed for growth on SD medium. Mutants displaying a significant growth defect were excluded at this stage. EV samples were then prepared from all remaining mutant strains using the quick protocol described previouslyl^62^, in batches of 20, with each batch including the reference strain KN99α. EV concentration was measured using NanoFCM. Mutants producing less than 0.7-fold or more than 1.3-fold the EV levels of the WT strain were subsequently re-tested independently using the same procedure. Only mutants for which the second measurement confirmed the initial result were retained. More than 60 mutants were tested at least three times in this study or in the context of other ongoing studies in the laboratory (Table S2).

### Colony forming units (CFU) assay and cell diameter measurements

Cells were grown under EV-producing conditions on SD agar plates for 24 h at 30°C, gently collected with a sterile loop, and resuspended in 0.22-µm-filtered 1× PBS. For viability assessment, cell numbers were determined using a Coulter Counter (Z Series, Beckman Coulter), and the suspension was diluted in sterile water to 1 × 10³ cells/mL. Aliquots of 200 µL were plated onto YPD agar plates and incubated for 48 h at RT, after which CFUs were determined. For cell diameter measurements, cells were imaged using an Olympus IX83 microscope equipped with a Hamamatsu ORCA-Flash cooled CCD camera and cellSens imaging software. Cell diameter was measured using ImageJ (n = 150).

### Glycerol-3-phosphate quantification

Cells were grown under EV-producing conditions on SD agar plates for 24 h at 30°C, gently collected with a sterile loop, and resuspended in sterile water. A total of 2 × 10⁸ cells in 500 µL of water were mixed with 400 µL of 425–600 µm acid-washed glass beads (Sigma, G8772-500G) and disrupted by bead beating for three 45-s cycles at 6.5 m/s using a FastPrep™ instrument. Samples were cooled on ice for 2 min and centrifuged for 10 min at 15,000 × g at 4°C. The supernatants were collected and transferred to fresh tubes for total glycerol and glycerol-3-phosphate quantification. Glycerol-3-phosphate was quantified by high-performance anion-exchange chromatography (HPAEC) using a Dionex ICS-3000 system equipped with a CarboPac PA1 column (250 × 4.6 mm; Thermo Scientific) and glycerol-3-phosphate (Ref: GC36159 GlpBio Technology Inc) as the standard. Samples were eluted at a flow rate of 1 mL/min with 18 mM NaOH for 15 min, followed by a linear gradient to 300 mM sodium acetate (AcONa) in 100 mM NaOH over 20 min and an isocratic step at 300 mM AcONa in 100 mM NaOH for 5 min. The column was equilibrated with the initial mobile phase for 20 min before each injection. Detection was performed using a pulsed electrochemical detector.

### Lipid extraction and UPLC-MS/MS analysis

Strains were grown under EV-producing conditions on SD agar plates for 18 h at 30°C, as previously described. Cells were gently collected in 0.22-µm-filtered PBS using a sterile loop and centrifuged at 5,000 × g for 15 min. The resulting pellet was resuspended in 8 mL of PBS and centrifuged at 15,700 × g for 10 min. The supernatant was discarded, and the cell pellet was flash-frozen in liquid nitrogen. Samples were stored at −80°C until shipment for lipidomic analysis (Metware Biotechnology Inc.USA).

Ultra-performance liquid chromatography-tandem mass spectrometry (UPLC-MS/MS) analysis was performed by MetwareBio (Woburn, MA, USA). Lipids were extracted by adding methyl tert-butyl ether (MTBE):methanol (3:1, v/v) containing an internal standard mixture. Samples were mixed for 15 min, followed by the addition of ultrapure water and vortexing for 1 min. Samples were then centrifuged at 12,000 rpm for 10 min, and the upper organic phase was collected and evaporated using a vacuum concentrator. The dried extracts were reconstituted in acetonitrile:isopropanol (1:1, v/v) prior to LC-MS/MS analysis. UPLC separation was performed using a Nexera LC-40 system (Shimadzu, Kyoto, Japan) equipped with an Accucore™ C30 column (2.6 µm, 2.1 × 100 mm; Thermo Fisher Scientific, Waltham, MA, USA) maintained at 45°C. The flow rate was 0.35 mL/min, and the injection volume was 2 µL. Mobile phase A consisted of acetonitrile/water (60:40, v/v) containing 0.1% formic acid and 10 mmol/L ammonium formate, while mobile phase B consisted of acetonitrile/isopropanol (10:90, v/v) containing 0.1% formic acid and 10 mmol/L ammonium formate. The gradient was as follows: 80:20 (A:B) at 0 min, 70:30 at 2 min, 40:60 at 4 min, 15:85 at 9 min, 10:90 at 14 min, 5:95 at 15.5 min, 5:95 at 17.3 min, 80:20 at 17.5 min, and 80:20 at 20 min.For the mass spectrometry linear ion trap (LIT) and triple quadrupole (QQQ) scans were acquired on a triple quadrupole-linear ion trap mass spectrometer QTRAP® 6500+ LC-MS/MS System (SCIEX, Framingham, MA, USA), equipped with an ESI Turbo Ion-Spray interface, operating in positive and negative ion mode and controlled by Analyst 1.6.3 software (SCIEX, Framingham, MA, USA). The electrospray ionization (ESI) source operation parameters were as follows: source temperature 500 ℃; ion spray voltage (IS) 5500 V(Positive),-4500 V(Negative); Ion source gas 1 (GS1), gas 2 (GS2), curtain gas (CUR) were set at 45, 55, and 35 psi, respectively. Instrument tuning and mass calibration were performed with polypropylene glycol solutions in QQQ mode (10 μmol/L) and LIT mode (100 μmol/L). QQQ scans were acquired using multiple reaction monitoring (MRM), with nitrogen as the collision gas at 5 psi. Declustering potential (DP) and collision energy (CE) were initially determined for each MRM transition and subsequently optimized. Specific sets of MRM transitions were monitored during each acquisition period according to the lipids expected to elute within the corresponding time interval. A quality control (QC) sample was prepared by pooling aliquots of all sample extracts to assess the reproducibility of the lipidomics workflow. During data acquisition, QC samples were analyzed approximately every 10 experimental samples, with a total of three QC injections.

### Statistical analysis

All statistical analyses were performed using GraphPad Prism 9 software (GraphPad Software Inc.). Data sets were tested for normal distribution using Shapiro–Wilk or Kolmogorov–Smirnov normality tests. In the cases in which the data passed the normality test, they were further analyzed using the unpaired Student’s t test or ordinary one-way analysis of variance. When at least one data set was non-normally distributed, we used the nonparametric Kolmogorov–Smirnov or Kruskal–Wallis test.

## Availability and accession number

Raw data are available at Bioprojects: PRJNA1540463, PRJNA928602 and GEO GSE133695

## Acknowledgments

We would like to thank Thierry Fontaine (Institut Pasteur) for his assistance with the glycerol-3-phosphate concentration measurements. The construction of the *C. neoformans* mutant collection in the Madhani laboratory (University of California, San Francisco), which was used in this study, was supported by NIH grant R01AI100272. Extracellular related work in G.J. laboratory is supported by the ANR grants ResistEV AAPG2021 CE35, FEVCOM AAPG2023 CE35, FunWallEV AAPG 2024 CE35, the PTR 2024 grant NanoEVs n°1785, and the CAPES COFECUB grant n**°** 52812YE. G.J is also a member the Pasteur International Unit “Fungal extracellular Vesicles”. AP was supported by the Pasteur-Roux-Cantarini fellowship of Institut Pasteur. JR was support by Fundação Carlos Chagas Filho de Amparo à Pesquisa do Estado do Rio de Janeiro - FAPERJ (E-26/203.607/2024, E-26/204.398/2025, E-26/210.568/2025, E-26/210.687/2025), Conselho Nacional de Desenvolvimento Científico e Tecnológico – CNPq (404273/2025-4), Inova FIOCRUZ - UFRJ 01/2025 program, and by the IDOR Pioneer Science Initiative (www.pioneerscience.org).

## Supplementary material

**Figure S1. EV production by 50 randomly chosen mutant strains. A.** EV quantification of knockout mutants; EV were isolated using the quick protocol methodology and measurement of particles were made using a VideoDrop instrument. **B.** FLC susceptibility spot assay. Serial dilutions of cells were spotted onto YPD medium containing fluconazole (15 µg/mL). Plates were incubated for 72 h at 30°C before imaging.

**Figure S2: Complementation of *vps25Δ* mutation restores EV production to WT level.** EV release by WT, *vps25Δ* and *vps25Δ +VPS25* mutants measured by nanoFCM after 24h of growth on SD plates. Data represent mean ± SD *** *p* < 0.001, one-way ANOVA followed by Dunnett’s test versus WT.

**Figure S3. The *had1Δ* mutant presents a slight growth defect on SD medium. Yet, extended growth does not suppress its EV production phenotype.**

**A.** Growth quantification based on total cell counts recovered per SD plate at 30°C. Data represent means ± SD. *p* < 0.001 (ANOVA followed by Tukey’s multiple comparisons test).

**B.** Quantification of particle release by WT and *had1Δ* strains using the quick extraction protocol after 28 or 30 h of incubation on SD plates, measured by nanoFCM. Data represent means ± SD *p* < 0.001 (two-tailed Student’s *t*-test).

**Figure S4. The *calcineurin pathway* mutants all present a slight growth defect on SD medium. Yet, extended growth does not suppress their EV production phenotype.**

**A,** Growth assessment by counting colonies recovered from SD plates after 24 h at 30 °C. Data represent mean ± SD (*** *P* < 0.001, one-way ANOVA followed by Tukey’s multiple-comparisons test).

**B**. EV release by WT, *cna1Δ*, *cnb1Δ* and *crz1Δ* mutants measured by nanoFCM after 28 and 30 h of growth on SD plates. Data represent mean ± SD (** *P* < 0.01; *** *P* < 0.001, one-way ANOVA followed by Dunnett’s test versus WT).

**Figure S5.** Exogenous glycerol does not suppress *had1Δ* phenotypes. *Left*: Quantification of EV release by WT and *had1Δ* strains grown in the presence or absence of 0.1 M glycerol, measured by NanoFCM (****P < 0.0001, one-way ANOVA followed by Dunnett’s multiple-comparisons test). *Right*: Heat map of fluconazole MIC values in the presence of 0.1 M glycerol; the white line indicates the MIC₅₀ (n = 6).

**Table S1. Time course RNA-seq analysis** A. lists of the genes associated with expression clusters during EV production in *C. neoformans* B. Manual functional annotation of C14 genes

**Table S2. Production of EVs by the 287 mutant strains considered in this study**

**Table S3. EV production by randomly chosen mutant strains.**

**Table S4: Viability of EV production mutants as evaluated by CFUs**

**Table S5. Lipid composition of WT vs Had1**

**Table S6: Plasmids used in this study**

**Table S7: Primers used in this study**

**Table S8: Strains used in this study**

## Notes

### Competing Interest Statement

The authors have declared no competing interest.

